# Association of CD44^−^/CD24^−^ Breast Cancer Cells with Late Stage Tumor Recurrence

**DOI:** 10.1101/2020.12.07.414599

**Authors:** Xinbo Qiao, Yixiao Zhang, Lisha Sun, Qingtian Ma, Jie Yang, Liping Ai, Jinqi Xue, Guanglei Chen, Hao Zhang, Ce Ji, Xi Gu, Haixin Lei, Yongliang Yang, Caigang Liu

## Abstract

Tumor metastasis remains the main cause of breast cancer-related deaths, especially the later breast cancer distant metastasis. This study assessed CD44^−^/CD24^−^ tumor cells in 576 tissue specimens for associations with clinicopathological features and metastasis and then investigated the underlying molecular events. The data showed that level of CD44^−^/CD24^−^ cells was associated with later postoperative distant tumor metastasis. Furthermore, CD44^−^/CD24^−^ triple negative cells could spontaneously convert into CD44^+^/CD24^−^ cancer stem cells (CSCs) with properties similar to CD44^+^/CD24^−^ CSCs from parental MDA-MB-231 cells in terms of gene expression, tumor cell xenograft formation, and lung metastasis *in vitro* and *in vivo*. Single-cell RNA sequencing identified RHBDL2 as a regulator that enhanced spontaneous CD44^+^/CD24^−^ CSC conversion, whereas knockdown of RHBDL2 expression inhibited YAP/NF-κB signaling and blocked spontaneous CD44^−^/CD24^−^ cell conversion to CSCs. These data suggested that the level of CD44^−^/CD24^−^ tumor cells could predict breast cancer prognosis, metastasis, and response to adjuvant therapy.

## Introduction

Breast cancer is the most prevalent malignancy in women, and its incidence is increasing worldwide, especially in developed countries (*Bianchini et al., 2016*; *Siegel et al., 2020*). Different treatment strategies, such as surgical resection, hormone therapy, targeted therapy, radiation therapy, and chemotherapy, have greatly improved the survival of breast cancer patients (*Bray et al., 2018*; *Burstein et al., 2014*; *Early Breast Cancer Trialists’ Collaborative, 2015; Khan et al., 2012*; *Saini et al., 2012*). However, as such advancements prolong patients’ survival, they also lead to an unfortunate and considerable number of patients facing the risk in developing later breast cancer metastasis, even 20–40 years after breast cancer diagnosis (*Sharma, 2018*). Notably, breast cancer metastasis occurring 5–8 years after initial surgical resection has become a significant cause of treatment relapse, tumor progression, and poor survival of patients (*Nishimura et al., 2013*); thus, further research on the underlying molecular mechanisms and gene alterations could help us to identify novel biomarkers and treatment strategies to effectively control breast cancer metastasis and progression. To date, clinical strategies for breast cancer treatment remain suboptimal. Currently, the best option available is continuous tamoxifen treatment for 10 years, which has been shown to reduce cancer recurrence and mortality of patients; however, a significant proportion of patients may be overtreated (*Bianchini et al., 2016*; *O’Conor et al., 2018*). In this regard, we also urgently need to discover novel biomarkers to predict treatment effectiveness and to improve treatment success and prognosis among breast cancer patients.

Indeed, tumor metastasis is a multistep process, involving tumor cell escape from the primary site, migration into neighboring tissues, extravasation, survival, and colonization, all leading to the formation of new tumor lesions at a secondary site (*Drabsch and ten Dijke, 2011; Klein, 2008; Scott et al., 2012*; *Syn et al., 2016*). The rate-limiting step of cancer metastasis is tumor growth at the secondary site, because the tumor lesion initially lacks sufficient vasculature to provide nutrients for cancer cell growth. Thus, the newly arrived tumor cells may grow to a certain size in the new and harsh microenvironment and then experience growth arrest in that organ. However, once they regain their proliferative ability, late metastasis will occur (*Langley and Fidler, 2007*). Molecularly, CD44+/CD24− tumor cells from the breast primary tumor lesion are associated with distance metastasis (*Abraham et al., 2005*), and these cells display increased motility and invasiveness (*Liu et al., 2010*) similar to chemoresistant cancer stem cells (CSCs) (*Velasco-Velazquez et al., 2011*). Previous studies have shown that CD44^+^/CD24^−^ breast CSCs might be a dominant factor in relapse of triple negative breast cancer (TNBC), due to their possession of potent self-renewal and differentiation capacities to differentiate into mature CD44^−^/CD24^−^, CD44^+^/CD24^+^, and CD44^−^/CD24^+^ cancer cells (*Geng et al., 2014*; *Wang et al., 2014*). Indeed, injection of up to 1000 breast CSCs was able to generate a solid tumor mass in immunocompromised mice (*Chaffer et al., 2011*; *Iliopoulos et al., 2011*). Thus, the number of breast CSCs in the secondary site could affect the efficiency of early metastasis formation, and stem cells are more prone to be resistant to chemotherapy (*De Angelis et al., 2019*). Moreover, previous studies reported that non-CSCs, such as CD44^−^/CD24^−^ TNBC cells, are able to spontaneously convert into CSCs to renew the CSC pool, resulting in chemoresistance(*Gruber et al., 2016*; *Kim et al., 2015*; *Ye et al., 2018*). Thus, the dormant CD44^−^/CD24^−^ tumor cells that have previously been colonized in the metastatic site may be able to spontaneously convert to CSCs to regain proliferative ability and drug resistance, resulting in later tumor metastasis. Therefore, detection of CD44^−^/CD24^−^ cells may be useful in the prediction of later breast cancer metastasis.

In the present study, we aimed to identify the molecular mechanisms by which CD44^−^/CD24^−^ cell to CSC conversion promotes later breast cancer metastasis. We first performed a retrospective analysis of CD44^−^/CD24^−^ breast cancer cells in tissue specimens from patients enrolled from three academic medical centers to identify any associations between the presence of these cells and postoperative tumor metastasis. We then performed *in vitro* and *in vivo* experiments to confirm CD44^−^/CD24^−^ cell conversion to CD44^+^/CD24^−^ CSCs and then assessed the properties of the converted CSCs in vitro and in vivo. This study provides novel insight into the role of CD44^−^/CD24^−^ tumor cells in later breast cancer metastasis and into the potential use of CD44^−^/CD24^−^ cells as a biomarker to predict survival and metastasis in breast cancer patients or targeting of RHBDL2 as a novel therapeutic approach in the future.

## Results

### Association of high CD44^−^/CD24^−^ tumor cell level with late breast cancer distant metastasis

CD44^+^/CD24^−^ breast CSCs were previously found to not have a significant impact on the prognosis of breast cancer patients (*Kaverina et al., 2017*; *Mylona et al., 2008*). In the present study, we performed a retrospective analysis of CD44^+^/CD24^−^ cell levels in 576 patients using immunofluorescence staining to examine their association with clinicopathological factors. The patients were divided into a training set (n= 355) and testing set (n= 221) (Table S1) for assessment of cell membrane proteins CD44 and CD24 (Fig. S1), leading to four types of breast cancer cells (CD44^−^/CD24_−_, CD44^+^/CD24^−^, CD44^−^/CD24^+^, and CD44^+^/CD24^+^), as shown in Fig. S2A. The level of CD44^−^/CD24^−^ cells was significantly higher in breast cancer tissues of patients with postoperative tumor metastasis in the training set (P<0.0001; Fig. 1A). The receiver-operating characteristic curve analysis showed a decision threshold of 19.5% CD44^−^/CD24^−^ cancer cells, and thus, we used this cut-off-point to perform a subgroup analysis (the discrimination criteria are shown in Fig. 1B). In the training set of patients, the metastasis rate was 1.97-fold higher in patients with a high level of CD44^−^/CD24^−^ breast cancer cells (>19.5%) than in those with a low level of CD44^−^/CD24^−^ tumor cells (<19.5%). Similar data were obtained after analysis of the three breast cancer subtypes (i.e., luminal: 63.1% vs. 32.58%; HER-2: 49.02% vs. 15%; TNBC: 56.96% vs. 20.28% for high vs. low CD44^−^/CD24^−^ tumor cells; Fig. 1C). Our univariate analysis showed that the CD44^−^/CD24^−^ cancer cell subgroup, CSCs percentage, lymph node metastasis, N stage, estrogen receptor and progesterone receptor status, and molecular subtype were all predictors of DFS (Table S2).

**Figure 1.**
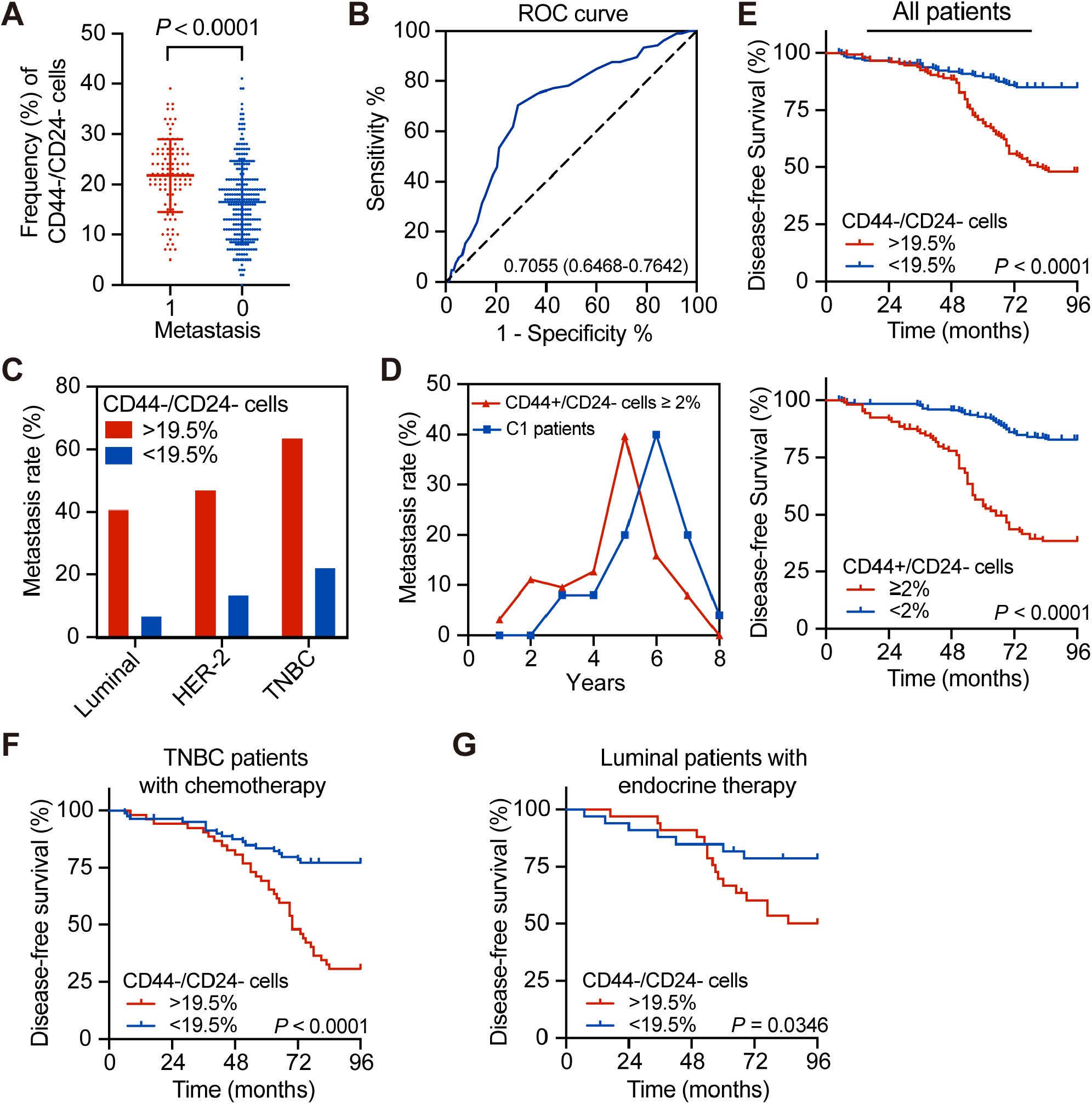
Association of high CD44^−^/CD24− cell percentage with later tumor distant metastasis. (A) Flow cytometry. Tissue samples from breast cancer patients with or without tumor metastasis were processed and subjected to flow cytometric analysis of CD44^−^/CD24^−^ cells. *P* values were determined by Student’s *t*-test. (B) Receiver-operating characteristic curve. The data in A were processed to generate the ROC curve to predict tumor metastasis events due to the percentage of CD44^−^/CD24^−^ cells. The area under the curve was 0.7055 (0.6468–0.7642; *P*<0.0001), and the threshold from the ROC curve analysis was 19.5%. (C) Metastasis rates in patients with different breast cancer molecular subtypes stratified by the percentage of CD44^−^/CD24^−^ cells. (D) Metastasis rates in patients with ≥2% CD44^+^/CD24^−^ cells (n=105) vs. C1 patients with <2% CD44^+^/CD24^−^ cells and ≥19.5% CD44^−^/CD24^−^ cells (n=69). (E) Kaplan–Meier curves. The DFS of breast cancer patients was plotted for Kaplan–Meier analysis and then subjected to the log rank test stratified by the percentage of CD44^−^/CD24^−^ cells vs. CD44^+^ cells. (F) Kaplan–Meier curves. The DFS of TNBC patients after chemotherapy was plotted for Kaplan-Meier analysis and then subjected to the log rank test stratified by ≥19.5% CD44^−^/CD24^−^ cells (n=52) vs. <19.5% CD44^−^/CD24^−^ cells (n=80). (G) Kaplan–Meier curves. The DFS of luminal breast cancer patients after endocrine therapy was plotted for Kaplan–Meier analysis and then subjected to the log rank test stratified by ≥19.5% CD44^−^/CD24^−^ cells (n=24) vs. <19.5% CD44^−^/CD24^−^ cells (n=23).

Furthermore, the DFS of patients with a high CD44^+^/CD24^−^ or CD44^−^/CD24^−^ tumor cell level was shorter than that of patients with a low frequency (Fig. 1E), which was also observed in the three breast cancer subtypes (Fig. S2D and S3A). The kinetic curve pattern analysis also showed that patients with a high CSC percentage had a high risk of developing early tumor metastasis within the first 3 years after diagnosis (Fig. S3C). Some patients with high CD44^−^/CD24^−^ tumor cell levels also possessed a high CSC percentage and showed a higher rate of postoperative metastasis. However, to exclude the effects of CSCs, we defined patients with <2% CD44^+^/CD24^−^ and > 19.5% CD44^−^/CD24^−^ tumor cells as the C1 group, and our data revealed that these C1 patients had a high risk of developing later tumor metastasis after 5–7 years (Fig. 1D).

### Level of CD44^−^/CD24^−^ breast cancer cells predicts success of postoperative TNBC treatment

We then confirmed our data using the testing set of patients and found that the metastasis rate was higher in patients with a high CD44^−^/CD24^−^ tumor cell level among all breast cancer subtypes (luminal: 63.1% vs. 32.58%; HER-2: 49.02% vs. 15%; TNBC: 56.96% vs. 20.28% for high vs. low CD44^−^/CD24^−^ cells; Fig. 2A). The DFS and OS of patients with a high CD44^−^/CD24^−^ percentage were shorter than those of patients with a low CD44^−^/CD24^−^ cell percentage (Fig. 2CD). Moreover, the DFS and OS of patients stratified by breast cancer subtype also were shorter with a high CD44^−^/CD24^−^ percentage vs. low CD44^−^/CD24^−^ percentage (Figure S5A,B). The kinetic curve pattern analysis also confirmed that C1 patients had a later metastasis peak than all other patients (Fig. 2B).

**Figure 2.**
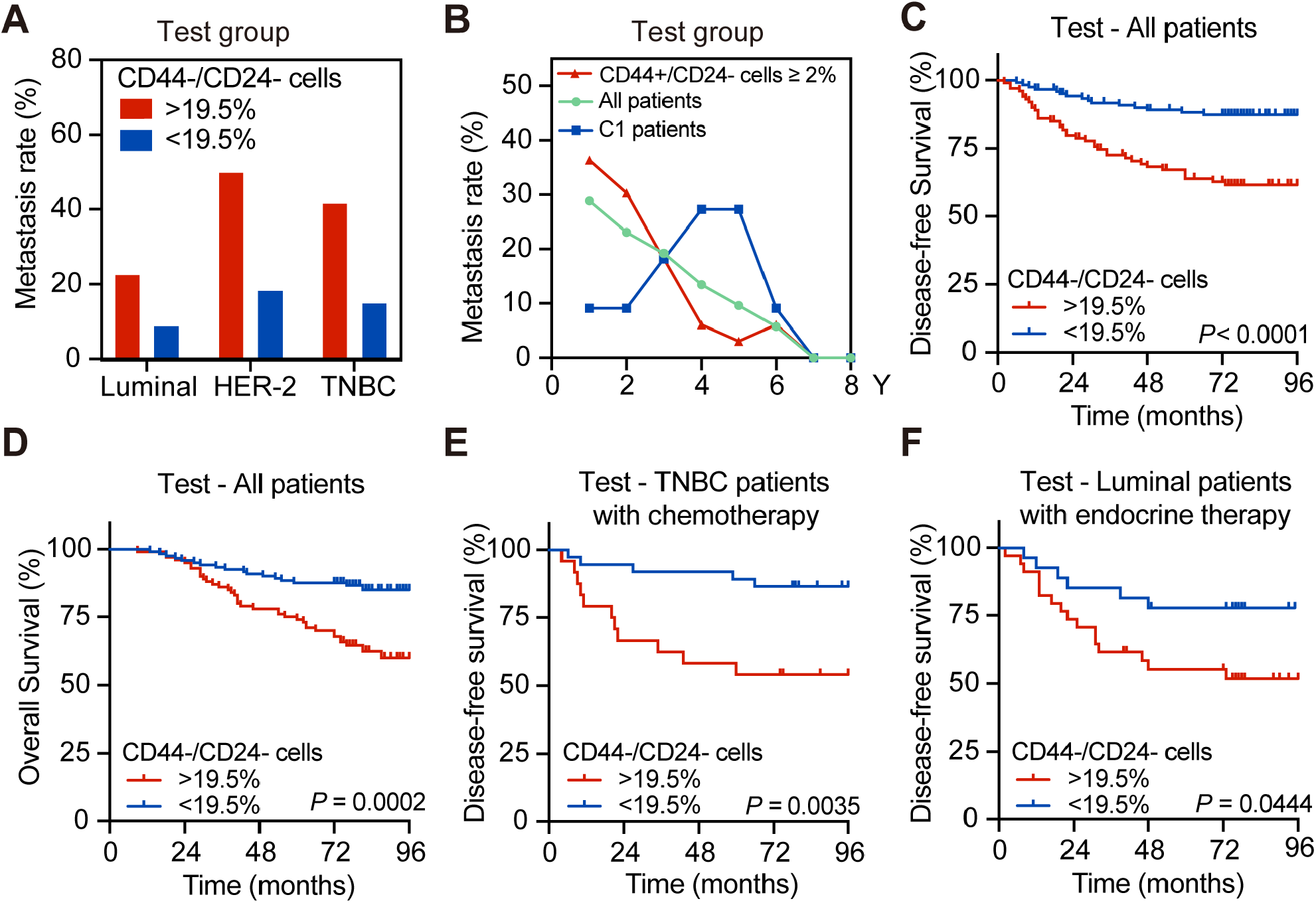
Ability of CD44^−^/CD24^−^ tumor cell percentage to predict the success of adjuvant TNBC treatment. (A) Postoperative metastasis rate in the test group of patients with different breast cancer molecular subtypes stratified by the percentage of CD44^−^/CD24^−^ cells. (B) Metastasis rate in all breast cancer patients (n=211) vs. patients with the ≥2% CD44^+^/CD24^−^ cells (n=68) vs. C1 patients with the <2% CD44^+^/CD24^−^ and ≥19.5% CD44^−^/CD24^−^ cells (n=153). (C and D) Kaplan–Meier curves. The DFS (C) and OS (D) of all patients in the testing group stratified by ≥19.5% CD44^−^/CD24^−^ (n=100) vs. <19.5% CD44^−^/CD24^−^ tumor cells (n=121). (E) Kaplan–Meier curves. The DFS in the test group of TNBC patients after chemotherapy stratified by ≥19.5% CD44^−^/CD24^−^ (n=24) vs. <19.5% CD44^−^/CD24^−^ tumor cells (n=35). (F) Kaplan–Meier curves. The DFS in the test group of luminal breast cancer patients after endocrine therapy stratified by ≥19.5% CD44^−^/CD24^−^ (n=22) vs. <19.5% CD44^−^/CD24^−^ tumor cells (n=19). *P*-values were obtained by log-rank test.

Furthermore, we analyzed the predictive value of the CD44^−^/CD24^−^ cell percentage for response to treatment. In the training set of patients, cases with a high CD44^−^/CD24^−^ cell level after chemotherapy had a higher risk of developing tumor metastasis 4 years after initial diagnosis and treatment than did cases with a low CD44^−^/CD24^−^ cell percentage (Fig. S3B). Moreover, a low CSC percentage led to different risks for developing tumor metastasis 5 years after diagnosis and adjuvant chemotherapy between the C1 and C0 patients (Fig. S3D). TNBC patients with high and low CD44^−^/CD24^−^ cell levels also had different risks of metastasis 4 years after diagnosis and chemotherapy, indicating that chemotherapy per se did not effectively inhibit tumor progression (metastasis) (Fig. 2F). In this regard, detection of CD44^−^/CD24^−^ cells in tumor tissues could be a valuable predictor of chemotherapy response. The data from our testing group of patients confirmed these results (Fig. 2E and S4CD). In addition, patients with luminal breast cancer treated with endocrine therapy also showed significantly different prognoses after stratification by percentage of CD44^−^/CD24^−^ cells in both the training and testing groups of patients (Fig. 1FG).

### Spontaneous conversion of CD44^−^/CD24^−^ TNBC cells into CD44^+^/CD24^−^ CSCs in vitro

Previous studies showed that non-CSCs are able to spontaneously convert into CSCs to renew the CSC pool, although it remained unclear whether the stem cells derived from the non-CSCs have the same biological behavior as the original stem cells (*Najafi et al., 2019*). In the present study, we first designed the experiments illustrated in Fig. 3A to characterize the percentages of different cell subtypes among MDA-MB-231 cells using cell culture and flow cytometric cell sorting of parental MDA-MB-468 cells. We found that the percentage of CD44^+^/CD24^−^ CSCs was 18.7% and that of CD44^−^/CD24^−^ CSCs was 73.4%. Next, we sorted the CD44^−^/CD24^−^ cells and continued to culture them in the stem cell (SC) and normal complete L15 medium for 7 days. We found that 28.3% of CD44^−^/CD24^−^ cells were able to spontaneously convert into CD44^+^/CD24^−^ CSCs, which was higher than the 24.7% observed when the cells were cultured in the normal complete L15 medium (Fig. 3B). Similar results were observed in the parental MDA-MB-468 cells (Fig. S5). Moreover, the percentage of CD44^+^/CD24^−^ CSCs was even higher after 21 days in culture than after 7 days in culture (Fig. 3C), suggesting that CD44^−^/CD24^−^ TNBC cells were able to spontaneously convert into CD44^+^/CD24^−^ CSCs.

**Figure 3.**
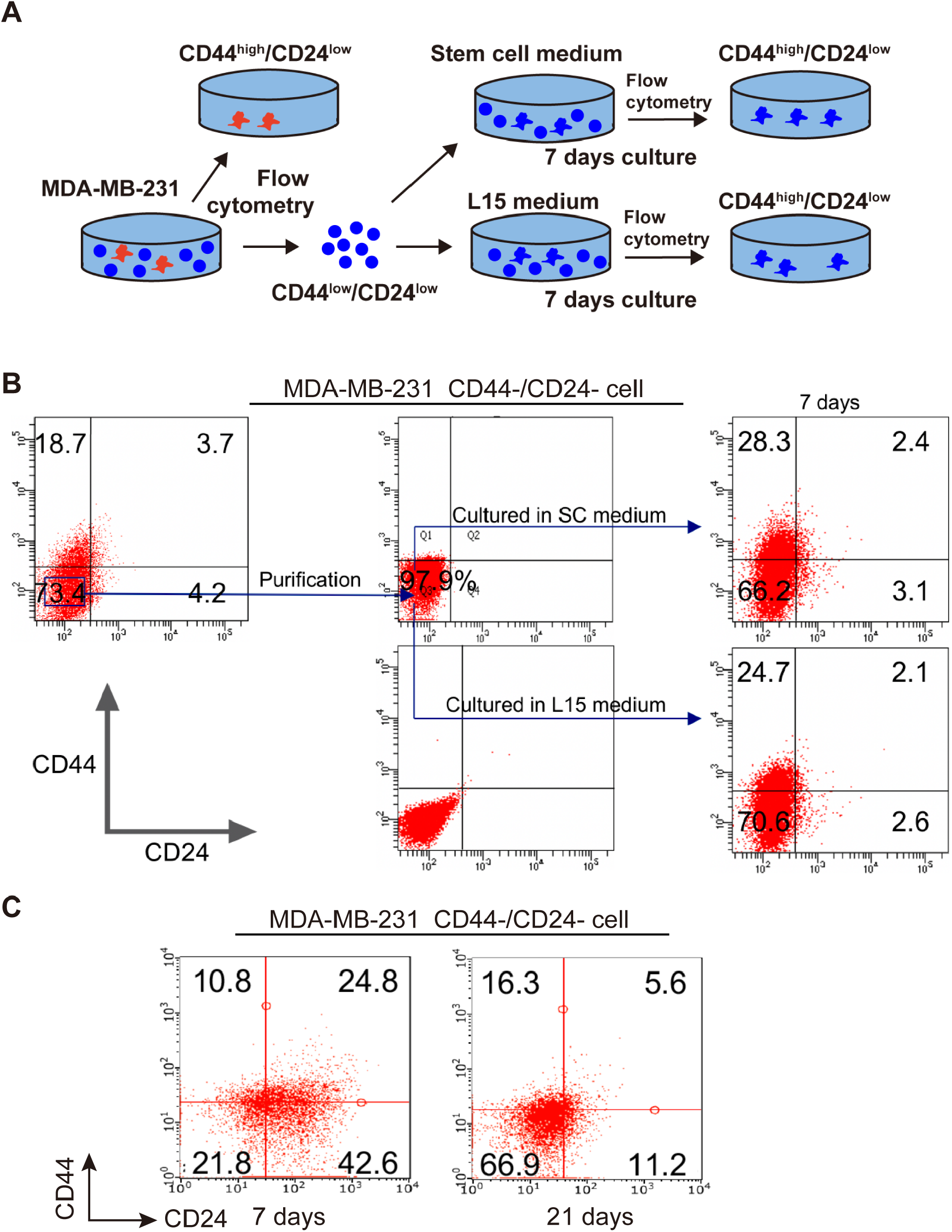
Spontaneous conversion of CD44^−^/CD24^−^ TNBC cells into CD44^+^/CD24^−^ CSCs. (A) Illustration of CD44^−^**/**CD24^−^ cell generation and sorting. (B) Flow cytometry. MDA-MB-231 cells were sorted with flow cytometry and then cultured in stem cell (SC) or L15 medium for 7 days. Then, the percentage of CD44^+^**/**CD24^−^ CSCs in CD44^−^**/**CD24^−^ MDA-MB-231 cells was analyzed by flow cytometry. (C) Flow cytometry. CD44^−^/CD24^−^ MDA-MB-231 cells were further sorted for CD44^+^/CD24^−^ CSCs by flow cytometry on days 7 and 21.

### Similar properties of newly converted CSCs from CD44^−^/CD24^−^ TNBC cells to those directly derived from TNBC cells in vitro

The clinical usefulness of CD44^−^/CD24^−^ cells in breast cancer lesions is obvious; thus, we explored the underlying mechanism by assessing whether these newly converted CD44^+^/CD24^−^ CSCs from CD44^−^/CD24^−^ MDA-MB-231 cells possessed comparable stemness properties to CSCs derived directly from parental MDA-MB-231 cells. We assessed changes in mammosphere formation, tumor cell differentiation ability, and expression of CSC biomarkers in the CSCs from these two sources in vitro as well as their tumorigenicity in vivo. We found that both CSC types, from the parental cell line and from their conversion from CD44^−^/CD24^−^ MDA-MB-231 cells, formed mammospheres similar in size and number (Fig. 4A) and displayed similar proliferative capacity (Fig. 4B). Culture of both types of CSCs for 7 days promoted their differentiation into different MDA-MB-231 cell subtypes at similar frequencies (Fig. 4CE). Western blot analysis showed that the expression of OCT4, SOX2, NANOG, NESTIN, and ABCB1 proteins was also comparable between these two types of CSCs (Fig. 4D). Thus, these results indicated that the newly converted CSCs from CD44^−^/CD24^−^ breast cancer cells and the CSCs derived directly from parental MDA-MB-231 cells exhibited comparable levels of tumor cell proliferation, differentiation, mammosphere formation, and stemness properties.

**Figure 4.**
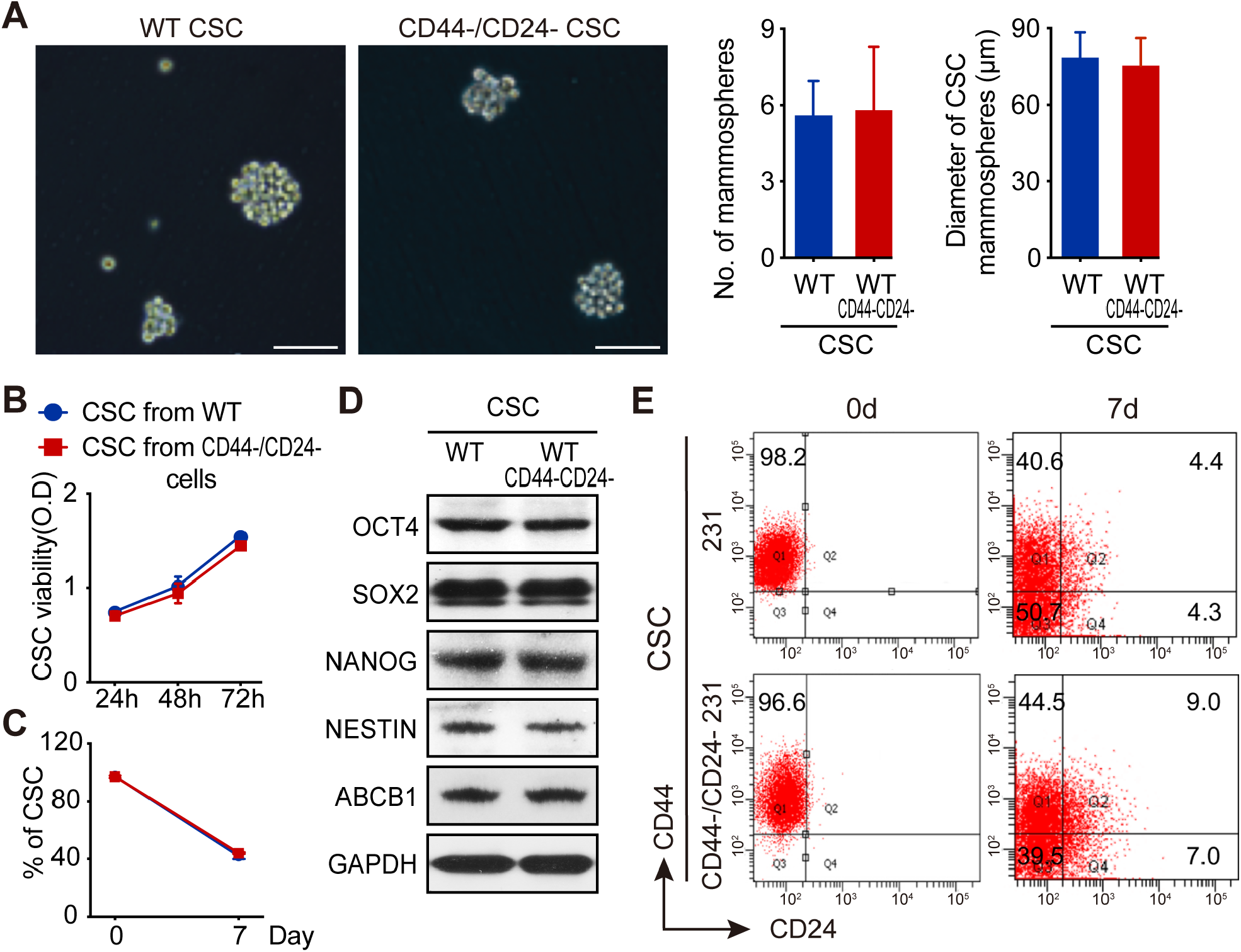
Similar properties of newly converted CSCs from CD44^−^/CD24^−^ TNBC cells to those of parental TNBC cells *in vitro*. (A) Morphology of CD44^+^/CD24^−^ CSCs derived from parental MDA-MB-231 (WT) and CD44^−^/CD24^−^ MDA-MB-231 cells. CD44^−^/CD24^−^ MDA-MB-231 cells were sorted by flow cytometry for CD44^−^/CD24^−^ cells, plated in cell culture dishes, and cultured for 7 days. Mammospheres were then photographed, and their number and size quantified. (B) Cell viability CCK-8 assay. The sorted CD44^+^/CD24^−^ CSCs (WT) and CD44^−^/CD24^−^ MDA-MB-231 cells from A were subjected to cell viability CCK-8 assay after culture for 24, 48, and 72 h. (C) Flow cytometry. The duplicated CSCs from WT and CD44^−^/CD24^−^ MDA-MB-231 cells were cultured in stem cell (SC) medium for 7 days and then sorted by flow cytometry. (D) Western blot. The sorted CD44^+^/CD24^−^ CSCs (WT) and CD44^−^/CD24^−^ MDA-MB-231 cells were then subjected to western blot analysis of the expression of stemness biomarkers, such as OCT4, SOX2, NANOG, Nestin, and ABCB1. (E) Flow cytometry. CD44^+^/CD24^−^ CSCs from parental MDA-MB-231 and CD44^−^/CD24^−^ cells converted MDA-MB-231 cells were cultured in stem cell (SC) medium for 7 days and then subjected to flow cytometry.

### Similar tumorigenesis and distant metastasis properties of CSCs derived from conversion of CD44^−^/CD24^−^ TNBC cells and directly from parental TNBC MDA-MB-231 cells in vivo

Next, we assessed whether the two types of CSCs functioned similarly in vivo. We inoculated BALB/c nude mice with equal numbers of the two types of CSCs, and 21 days later, we observed xenograft tumors of similar size and weight (Fig. 5A). Our flow cytometric analysis of these xenograft tumor cells showed similar frequencies of different breast cancer cell subtypes (Fig. 5B). Furthermore, injection of equal numbers of the two types of CSCs into the left ventricle of NOD/SCID mice also resulted in a similar incidence of tumor cell lung metastasis and metastasized tumors of similar size and weight (Fig. 5C). H&E staining of the metastasized tumors showed they had similar histology (Fig. 5C). Thus, the CSCs obtained from CD44^−^/CD24^−^ TNBC cells had similar abilities of tumorigenesis and metastasis in vivo compared with those obtained directly from parental TNBC cells.

**Figure 5.**
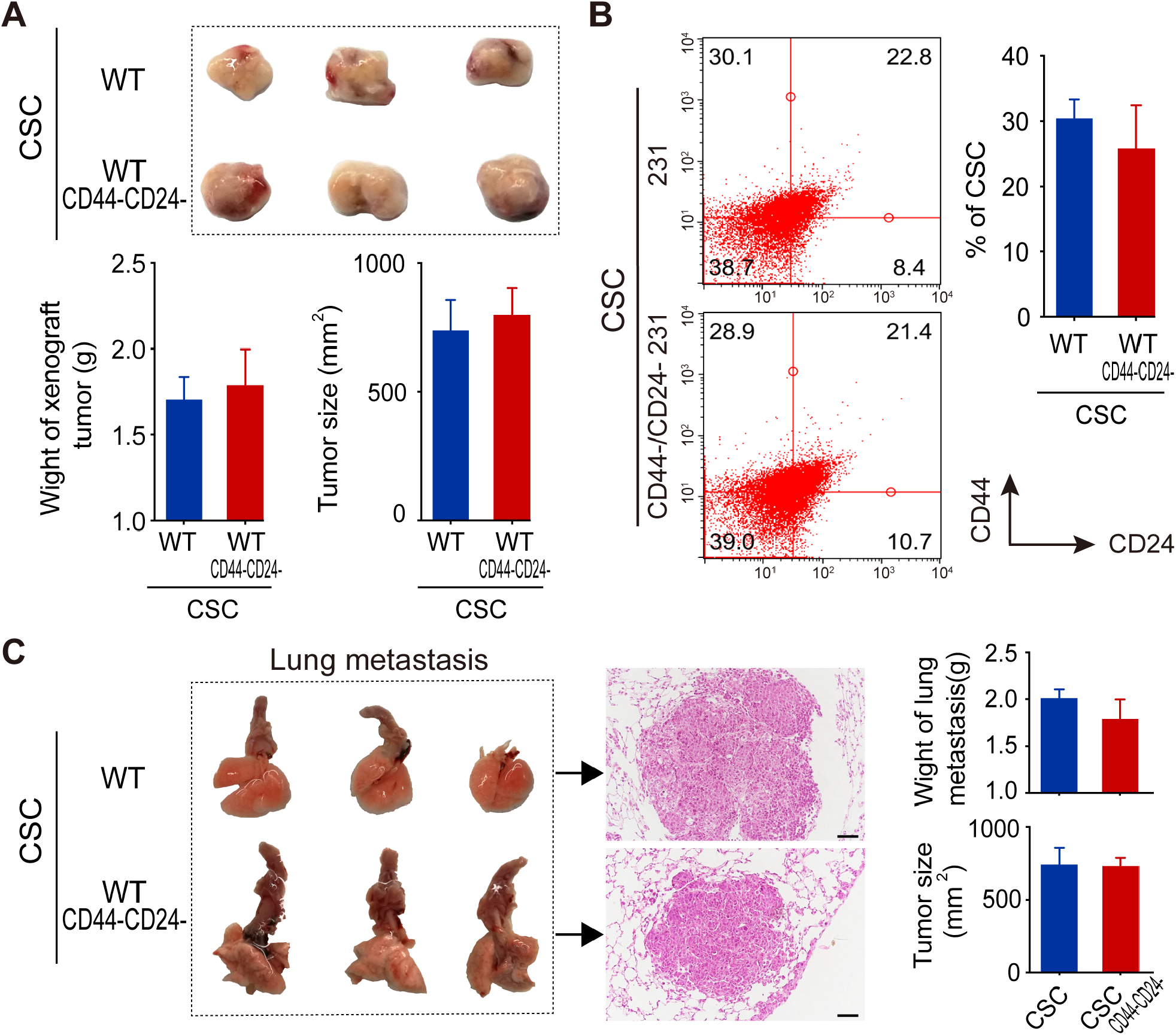
Similar properties of newly converted CSCs from CD44^−^/CD24^−^ TNBC cells to those of parental TNBC cells *in vivo*. (A) Tumor cell xenografts. Equal numbers of CSCs from CD44^+^/CD24^−^ (WT) and CD44^−^/CD24^−^ MDA-MB-231 cells were injected into BALB/c nude mice and grown for 21 days. After that, the mice were sacrificed, and the tumor xenografts were harvested, photographed (n=5-8 per group), and weighed. Tumor size also was measured. The data in the graphs for tumor weight and size are mean ± standard deviation (SD) for each group from three independent experiments. \*\**P*<0.01 using Student’s *t*-test. (B) Flow cytometry. The percentages of CSCs in different subtypes of MDA-MB-231 cells from individual xenografted tumors in A were analyzed by flow cytometry. (C) In vivo tumor cell lung metastasis. Equal numbers of CSCs from CD44^+^/CD24^−^ and CD44^−^/CD24^−^ converted MDA-MB-231 cells were injected into the tail vein of NOD/SCID mice and grown for 21 days to assess their lung metastasis. After that, the mice were sacrificed, and the tumor cell lung metastasis was resected. The tissue processed was for hematoxylin and eosin (H&E) staining and photographed (n=5–8 per group), and tumor weight and size were measured in each group.

### Involvement of RHBDL2 in spontaneous CD44^−^/CD24^−^ tumor cell conversion to CD44^+^/CD24^−^ CSCs

To further understand the molecular mechanisms underlying the CD44^−^/CD24^−^ cell conversion of TNBC cells, we first sorted CD44^−^/CD24^−^ cells from parental MDA-MB-231 cells and transfected them with an OCT4-EGFP plasmid (indicative of stem-like cells; Fig. 6A). We then isolated and cultured the single cells in complete L15 medium and performed RNA-seq to identify DEGs at 24 or 72 h after gene transfection. Our analysis revealed 11 DEGs associated with tumor cell differentiation and dedifferentiation after 24 h in cell culture (including RHBDL2, DSCC1, ZNF710, ATP8B3, and others; Fig. 6B and Fig. S6A). Our GO and KEGG analyses revealed that the reverse differentiation predominantly affected the expression levels of proteins involved in cell organelle formation, metabolism, signal transduction, and transcriptional regulation (Fig. S6BC). Furthermore, we performed Kaplan-Meier curve analysis and log-rank test to analyze associations between select DEGs and breast cancer prognosis using The Cancer Genome Atlas (TCGA) data (Fig. 6C). This analysis showed that high expression of HIST1H4H, ZNF710, RHBDL2, DSCC1, ARL6IP1, and PPME1 mRNAs was associated with a poor prognosis in breast cancer patients, whereas low expression of SLC35F5, G2E3, ATP8B3, and MED22 mRNAs also was associated with a poor prognosis of breast cancer patients. Furthermore, we cultured breast cancer cells for 7 days and then divided them into CD44^+^/CD24^−^ breast cancer stem cells, CD44^−^/CD24^−^ double negative cells, and dedifferentiated stem cells for detection of the expression levels of different mRNAs. We found that RHBDL2 expression was high during tumor cell dedifferentiation, suggesting that RHBDL2 may play a key role in the process of breast cancer cell dedifferentiation.

**Figure 6.**
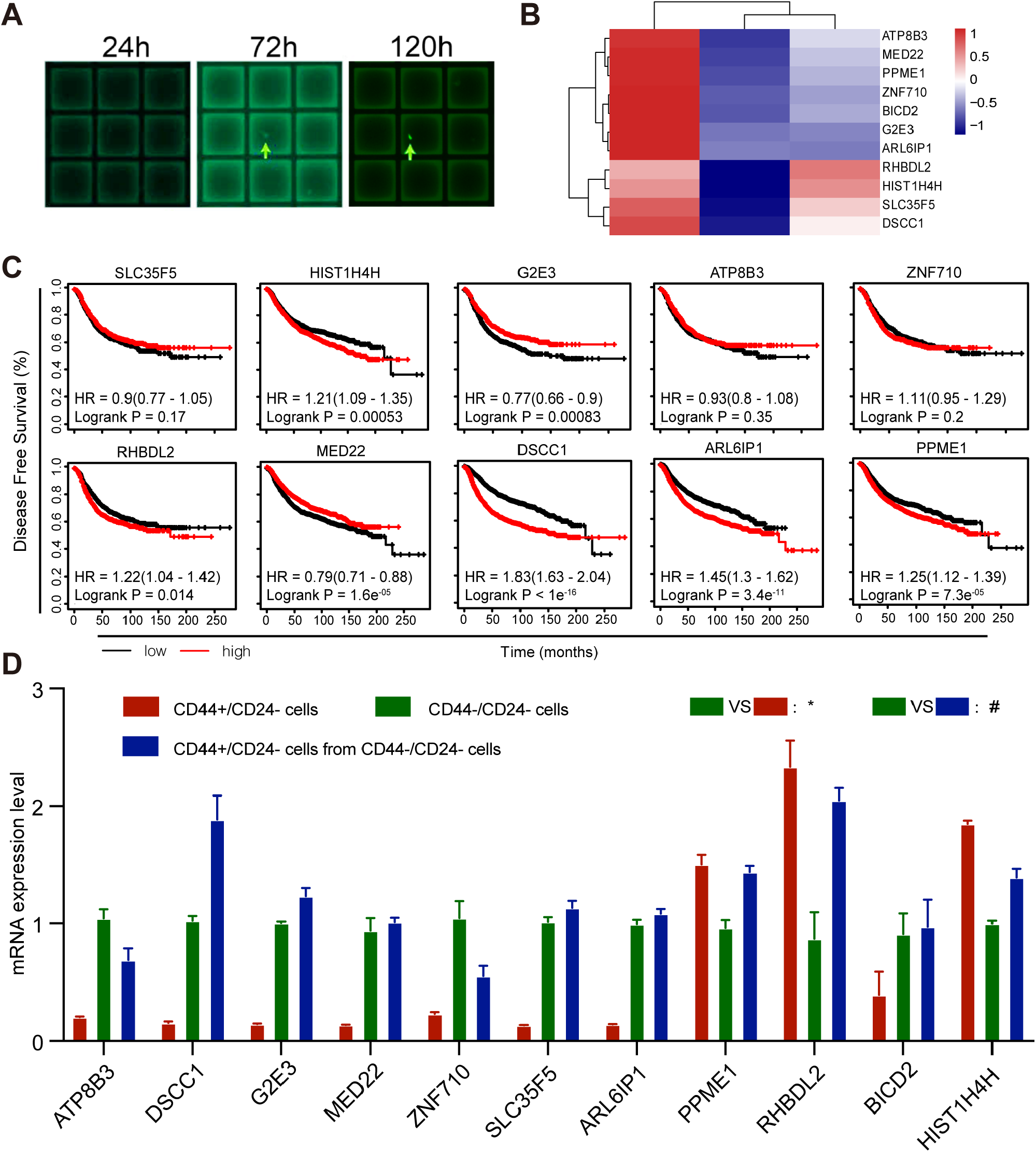
RHBDL2 involvement in the regulation of spontaneous conversion of CD44^−^/CD24^−^ MDA-MB-231 cells into CD44^+^/CD24^−^CSCs as analyzed by single-cell RNA-seq. (A) Morphology. The CD44^−^/CD24^−^ MDA-MB-231 cells were grown and transfected with OCT4-EGFP plasmid to illustrate the stem-like cells, cultured for 24 h, 72 h, and 120 h, and photographed. RNA-seq analysis. The CD44^−^/CD24^−^ MDA-MB-231 cells were grown and sorted by flow cytometry, and the resultant cells were grown and transfected with pGL3-OCT4-EGFP. Single cells were then captured at 24, 72, and 120 h, isolated, and subjected to RNA-seq analysis of DEGs. The major DEGs are shown in a heat map. (C) Kaplan–Meier plot for survival analysis. We generated Kaplan-Meier curves and applied the log-rank test to analyze associations of some selected DEGs with breast cancer prognosis using The Cancer Genome Atlas (TCGA) data according to the data in B. (D) qRT-PCR. The expression levels of different mRNAs selected from B were analyzed using qRT-PCR in CD44^+^/CD24^−^, CD44^−^/CD24^−^, and CD44^+^/CD24^−^ cells derived from CD44^−^/CD24^−^ cells after 7 days in culture.

### Downregulated RHBDL2 expression inhibits YAP/nuclear factor (NF)-κB/interleukin (IL)-6 signaling and spontaneous CD44^−^/CD24^−^ cell conversion to CD44^+^/CD24^−^ CSCs

To further explore how RHBDL2 and the related gene signaling are involved in the control of spontaneous CD44^−^/CD24^−^ cell conversion to CD44^+^/CD24^−^ CSCs, we first assayed the levels of total YAP1 and phosphorylated YAP1 in MDA-MB-231 cells (Fig. 7A) and found that phosphorylated YAP1 was increased with RHBDL2 silencing. Previous studies revealed that the YAP signaling is able to suppress USP31 expression, a potent inhibitor of NF-κB signaling in cells (17, 20). We thus further examined USP31 expression and NF-κB phosphorylation in CD44^−^/CD24^−^ cells sorted from both the negative control (NC) and RHBDL2-silenced MDA-MB-231 cells. Our results showed that USP31 expression was significantly higher in CD44^−^/CD24^−^ RHBDL2-silenced cells than in NC MDA-MB-231 cells, whereas the level of phosphorylated NF-κB was significantly lower in CD44^−^/CD24^−^ RHBDL2-silenced cells than in CD44^−^/CD24^−^ cells from negative control siRNA-transfected MDA-MB-231 cells (Fig. 7B). Furthermore, we investigated changes in the YAP1 distribution in the nucleus and cytoplasm of CD44^−^/CD24^−^ RHBDL2-silenced cells vs. CD44^−^/CD24^−^ cells from negative control siRNA-transfected MDA-MB-231 cells. Our data showed that both the nuclear and cytoplasmic levels of YAP1 protein were lower in RHBDL2-silenced cells than in negative control siRNA-transfected cells (Fig. 7C).

**Figure 7.**
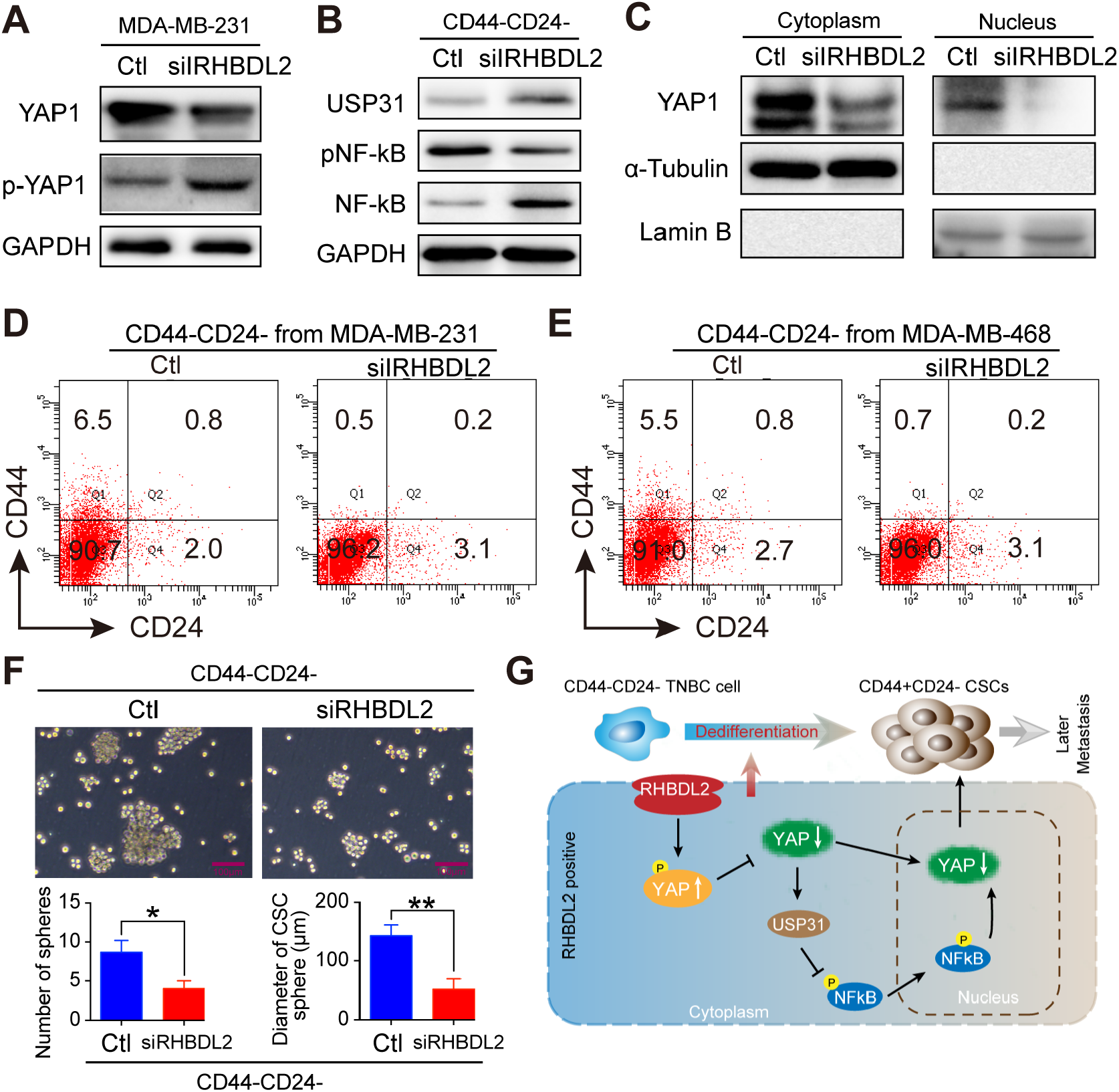
Mitigation of CD44^−^/CD24^−^ cell dedifferentiation into CSCs with RHBDL2 silencing in TNBC cells after inhibition of YAP1/NF-κB signaling. (A) YAP1 and phosphorylated YAP1 levels in MDA-MB-231 and RHBDL2-silenced MDA-MB-231 cells determined by western blot analysis. (B) USP31 and phosphorylated NF-κB levels in CD44^−^/CD24^−^ and RHBDL2-silenced CD44^−^/CD24^−^ cells determined by western blot analysis. (C) Cytoplasmic and nuclear YAP1 protein levels in MDA-MB-231 and RHBDL2-silenced MDA-MB-231 cells determined by western blot analysis. (D) Mammosphere formation by CSCs. MDA-MB-231 CD44^−^/CD24^−^ cells were grown and transfected with the negative control or RHBDL2-specific siRNA. After culture in SC medium for 7 days, mammospheres were randomly selected and photographed at 400×. The graphs show quantified data for mammospheres visible in five or more images from each of three wells. (E, F, and G) Illustration of the possible mechanisms underlying the enhanced conversion of CD44^−^/CD24^−^ cells into CD44^+^/CD24^−^ CSCs among RHBDL2-silenced TNBC cells.

We next selected the CD44^−^/CD24^−^ cells using flow cytometric sorting and transfected them with an RHBDL2-specific siRNA. After 7 days, the cell content was observed by flow cytometric analysis. The results showed that the ratio of CD44^+^/CD24^−^ CSCs dedifferentiated from NC cells was 6.5% and the ratio of siRHBDL2 cells converted to stem cells was 0.7% (Fig. 7D). Similar results were observed for parental MDA-MB-468 cells (Fig. 7E), with rates of CSC production of 5.5% (NC) and 0.5% (siRHBDL2). Then, we transfected CD44^−^/CD24^−^ MDA-MB-231 cells with negative control or RHBDL2-specific siRNA for 48 h (the transfection efficiency reached 62% and the knockdown of RHBDL2 expression was significant; P<0.01). After 7 days in culture, RHBDL2 silencing in CD44^−^/CD24^−^ MDA-MB-231 cells dramatically reduced the level of CD44^+^/CD24^−^ CSCs (P<0.01; Fig. 7F), and the number and size of the formed mammospheres were significantly less compared with those of CD44^−^/CD24^−^ cells from negative control-transfected MDA-MB-231 cells (P<0.05 and P<0.01, respectively; Fig. 7F). Specifically, RHBDL2 downregulated YAP1 expression to increase USP31 and decrease NF-κB signaling, and in turn, induce spontaneous conversion of CD44^−^/CD24^−^ TNBC cells into CD44^+^/CD24^−^ CSCs, whereas knockdown of RHBDL2 expression had the opposite effects in TNBC cells (Fig. 7G).

## Discussion

Our current study revealed that (a) CD44^−^/CD24^−^ TNBC MDA-MB-231 cells were able to spontaneously convert to CD44^+^/CD24^−^ CSCs, which is consistent with previous studies showing that non-tumor and neoplastic non-stem cells and CD44^−^/CD24^−^ breast cancer cells can spontaneously convert into CSCs *in vitro* (*Italiano and Shivdasani, 2003; Zoppino et al., 2010*); (b) CD44^+^/CD24^−^ CSCs converted from CD44^−^/CD24^−^ cells had similar biological properties compared to CSCs isolated from parental MDA-MB-231 cells in terms of proliferation capacity, gene expression, tumor cell xenograft formation, and lung metastasis *in vitro* and *in vivo*; (c) the percentage of CD44^−^/CD24^−^ cells was associated with later postoperative distant tumor metastasis; (d) the percentages of CD44^−^/CD24^−^ and CD44^+^/CD24^−^ cells, tumor N stage, and molecular subtypes were all predictors of DFS for breast cancer patients; and (e) mechanistically, knockdown of RHBDL2 expression inhibited YAP/NF-κB/IL-6 signaling and blocked the spontaneous CD44^−^/CD24^−^ cell conversion to CD44^+^/CD24^−^ CSCs. In conclusion, this study demonstrated that the CD44^−^/CD24^−^ cell population in breast cancer lesions was associated with breast cancer prognosis and distant metastasis as well as the success of adjuvant therapy. Further research is needed to investigate the role of RHBDL2 in breast cancer development and tumor cell dedifferentiation as a potential novel target in control of breast cancer progression.

The dedifferentiation (conversion) of non-CSC tumor cells into CSCs also occurs in other types of human cancers, such as glioblastoma and intestinal stroma melanoma, and contributes to intra- and inter-tumor heterogeneity (*Stepanova et al., 2003*; *Tzimas et al., 2006*; *Wei et al., 2019*). Although the underlying molecular mechanisms remain to be defined, CSC plasticity and heterogeneity could induce tumor progression and resistance to therapy (*Das et al., 2020*; *Kilmister et al., 2020*; *Martin-Castillo et al., 2013*; *Thankamony et al., 2020*). For example, intratumoral heterogeneity is a major ongoing challenge for effective cancer therapy, while CSCs are responsible for intratumoral heterogeneity, therapeutic resistance, and metastasis, which may be because cancer cells exhibit a high level of plasticity and an ability to dynamically switch between CSC and non-CSC states or among different subsets of CSCs (*Thankamony et al., 2020*). Another previous study showed that the differentiated non-CSCs are able to revert to trastuzumab-refractory, CS-like cells by activation of the tumor cell epithelial-to-mesenchymal transition process (*Martin-Castillo et al., 2013*). Conversion of non-CSCs to CSCs due to changes in tumor microenvironment and epigenetics could result in tumor progression and therapy failure (*Das et al., 2020*). Our current study further confirmed that the CD44^−^/CD24^−^ TNBC MDA-MB-231 cells spontaneously converted to CD44^+^/CD24^−^ CSCs and such CSCs possessed similar biological properties to CSCs isolated from parental MDA-MB-231 cells in terms of proliferation capacity, gene expression, tumor cell xenograft formation, and lung metastasis *in vitro* and *in vivo*. Indeed, the breast cancer MDA-MB-231 cell line was isolated and established from a pleural effusion of a 51-year-old white female in 1973 by Dr. R. Calleau of the University of Texas M.D. Anderson Cancer Center and does not express estrogen and progesterone receptors or HER2; thus, the MDA-MB-231 line is a TNBC cell line (*Cailleau et al., 1974*) (https://en.wikipedia.org/wiki/List_of_breast_cancer_cell_lines#MDA-MB-231) with naturally high CD44 expression and low CD24 expression (*Murohashi et al., 2010*). However, the factors that regulate this conversion of CD44^−^/CD24^−^ TNBC non-CSCs to CD44^+^/CD24^−^ CSCs remained incompletely understood, and our current study provides some insightful information, i.e., knockdown of RHBDL2 expression inhibited YAP/NF-κB/IL6 signaling and blocked the spontaneous CD44^−^/CD24^−^ cell conversion to CD44^+^/CD24^−^ CSCs. However, but further studies are needed to fully understand how RHBDL2 and the downstream YAP/NF-κB/IL-6 signaling mediate the regulation of this conversion.

Furthermore, our multicenter retrospective analysis of CD44^−^/CD24^−^ cancer cells in breast cancer tissue specimens showed that the CD44^−^/CD24^−^ cancer cell population was associated with postoperative breast cancer metastasis. Our multivariate Cox regression analysis revealed that percentages of CD44^−^/CD24^−^ and CD44^+^/CD24^−^ cells, tumor N stage, and molecular subtypes were all predictors of DFS for breast cancer patients. The percentage of CD44^−^/CD24^−^ cancer cells could be a better predictor of later breast cancer metastasis (up to 12 years after initial breast cancer diagnosis) than of early metastasis (within 5 years). Our C1 subgroup of patients had a higher risk of later metastasis at 5–7 years after surgery. The ROC curve analysis showed that the percentage of CD44^−^/CD24^−^ cells could predict later metastasis in cases with a low CSC percentage. These results further suggest that CD44^−^ /CD24^−^ cells could be able to spontaneously convert into CSCs and cause distant metastasis, although another theory of the clonal evolution also indicates that successive mutations accumulating in a given cell line may lead to clonal outgrowth in response to microenvironmental selection pressures (*Meacham and Morrison, 2013*).

In addition, our RNA-seq analysis identified several DEGs during the dynamic process of CD44^−^/CD24^−^ TNBC cell conversion into CSCs, i.e., expression of RHBDL2, YAP1, and other genes were significantly increased. Consistently, YAP/USP31/NF-κB signaling was activated in CD44^−^/CD24^−^ MDA-MB-231 cells, which is consistent with previous research showing that YAP1-mediated suppression of USP31 enhances NF-κB activity to promote sarcomagenesis (*Kemeny and Fisher, 2018; Mehta et al., 2018*). YAP1 can enhance CSC stemness in several types of human cancers, and aberrant YAP1 activation was associated with a low level of TNBC differentiation and poor survival of breast cancer patients (*Bora-Singhal et al., 2015*; *Hansen et al., 2015*). Our current data could indicate that upregulated RHBDL2 during CD44^−^/CD24^−^ TNBC cell conversion into CSCs could enhance YAP expression to, in turn, inhibit USP31 and attenuate its inhibition of NF-κB signaling, leading to the enhanced conversion of CD44^−^/CD24^−^ TNBC cells into CSCs.

In summary, postoperative breast cancer metastasis, especially later metastasis (e.g., ≥5 years after diagnosis), is an important unresolved issue. Our current results showed that patients with high CD44^−^/CD24^−^ cell populations had a high risk of developing metastasis 5–7 years after diagnosis. Knockdown of RHBDL2 expression caused YAP1 reduction and enhanced USP31/NF-κB signaling, leading to inhibition of the conversion of CD44^−^/CD24^−^ TNBC cells into CSCs. Furthermore, inhibition of the YAP1 signal could attenuate or even reverse the dedifferentiation process of TNBC cells. These findings provide novel insight into the molecular process of the dedifferentiation of non-CSCs into CSCs, which was been reported in glioblastoma, intestinal stroma melanoma, and other cancers, leading to cancer progression. Thus, our current findings suggest a novel postoperative therapeutic strategy for TNBC. Future studies will investigate targeting of RHBDL2 to control breast cancer resistance to chemotherapy.

## Materials and Methods

### Patients and tissue specimens

Paraffin-embedded cancer tissue samples were collected from 576 breast cancer patients who underwent breast cancer surgery between June 2005 and April 2013 in China Medical University Affiliated Hospital (Shenyang, Liaoning, China), Liaoning Cancer Hospital (Shenyang, Liaoning, China), and Dalian Municipal Central Hospital Affiliated of Dalian Medical University (Dalian, Liaoning, China). These patients were diagnosed with invasive breast cancer histologically according to the World Health Organization (WHO) breast cancer classifications, 4^th^ edition (*Tan et al., 2015*) and classified according to breast cancer TNM staging(*Li et al., 2012*). The patients’ clinicopathological data and follow-up data were collected from their medical records or via telephone interview. The inclusion criteria were: (a) surgical treatment for breast cancer; (b) complete information regarding clinicopathological characteristics; and (c) complete follow-up data. Disease-free survival (DFS) was defined as the time from the date of surgery to the date of distant metastasis, while overall survival (OS) was defined as the time from the date of surgery to the date of death. This study was approved by the ethics committee of all three hospital review boards, and each participant signed an informed consent form before being included in this study.

### Immunofluorescence staining

Paraffin-embedded cancer tissue blocks from 576 breast cancer patients were collected and used to construct a tissue microarray (*Edge and Compton, 2010*). The levels of CD44 and CD24 expression were assayed by using a double immunofluorescence staining in these tissue microarray sections. Briefly, 4-μm-thick sections of the tissue microarray were prepared, deparaffinized, rehydrated, and subjected to the antigen retrieval as described previously. Then the sections were incubated in 10% fetal bovine serum (FBS, Cellmax, Lanzhou, China) at room temperature for 1 h before incubation with a mouse anti-human CD44 monoclonal antibody (Cat. #3570; Cell Signaling Technology, Danvers, MA, USA) and a rabbit anti-human CD24 antibody (Cat. #ab202073; Abcam, Cambridge, MA, USA) at 4°C overnight. On the next day, the sections were washed with phosphate-buffered saline (PBS) briefly three times and further incubated with the Alexa Fluor 647-labeled rabbit goat anti-mouse IgG and Alexa Fluor 488-labeled goat anti-rabbit IgG, followed by nuclear staining with 4’,6-diamidino-2-phenylindole (DAPI). The fluorescence signals of the immunostained tissue sections were observed and recorded under a fluorescence microscope (Nikon, Tokyo, Japan). The percentages of CD44^+^/CD24^−^ CSCs and CD44^−^/CD24^−^ cells were calculated among a total of 2000 tumor cells from at least three sections in a blinded manner. The fluorescence images were obtained using a Nikon E800 upright microscope with NIS-Elements F3.0 (Nikon) and analyzed using ImageJ software (National Institute of Heath, Bethesda, MD, USA).

### Cell lines and culture

Human TNBC cell lines MDA-MB-231 and MDA-MB-468 were obtained from American Type Culture Collection (ATCC; Manassas, VA, USA) and maintained in Leibovitz’s L15 medium (Thermo Fisher, Carlsbad, CA, USA) supplemented with 10% FBS, 100 U/ml penicillin, and 100 μg/ml streptomycin (as the complete L15 medium) in a humidified incubator without addition of CO_2_ at 37°C.

The sorted CD44^−^/CD24^−^ cells (see below) were then cultured in the complete L15 medium or the stem cell (SC) medium (10% human MammoCult Proliferation Supplements in MammoCult Basal Medium, Stem Cell Technologies, San Diego, CA, USA) for 7 days. The cells were then subjected to different assays (see below).

### Flow cytometry

Parental MDA-MB-231 (wild-type, WT), RHBDL2-silenced MDA-MB-231, MDA-MB-468/Ctrl, and MDA-MB-468/sh cells were grown and subjected to incubation with fluorescein isothiocyanate (FITC)-conjugated anti-CD24 (Cat. #311104; BioLegend, San Diego, CA, USA) and PE-conjugated anti-CD44 (Cat. #338808; BioLegend) antibodies. Cells stained with an isotype control, FITC-anti-CD24 or PE-anti-CD44 alone served as negative controls. Then, the percentages of CD44^−^/CD24^−^, CD44^−^/CD24^+^, CD44^+^/CD24^+^, and CD44^+^/CD24^−^ cells were analyzed by using a FACSAria III flow cytometer (BD Biosciences, San Jose, CA, USA).

### siRNA and cell transfection

The yes-associated protein (YAP)-specific siRNA (5’-CAGUGUUUCAUACUCAAAU-3’), RHBDL2-specific siRNA (5’-CAUACUUGGAGAGAGAGCUAATT-3’), and negative control siRNA (5’-UUCUCCGAACGUGUCACGUTT-3’) were obtained from GenePharma (Shanghai, China) and used to knockdown expression of the corresponding genes. In particular, CD44^−^/CD24^−^ MDA-MB-231 cells were cultured in 6-well plates overnight at a density of 5 × 10^5^ cells/well until they reached 70% confluency when they were transiently transfected with the YAP-specific siRNA, RHBDL2-specific siRNA, or negative control siRNA using Mission siRNA transfection reagents (Sigma-Aldrich, St. Louis, MO, USA) for 48 h. The efficacy of silencing for each specific gene was evaluated by using western blot analysis.

### Western blot analysis

Cells were lysed in radioimmunoprecipitation (RIPA) lysis buffer containing phenylmethane sulfonyl fluoride (PMSF) and protease and phosphatase inhibitors. After centrifugation at 12000 rpm at 4°C for 20 min, the supernatants were collected, and the total protein concentration was measured using a Pierce BCA Protein Assay Kit (Thermo-Fisher) according to the manufacturer’s instructions. Next, these individual cell lysate samples (50 μg/lane) were separated by 12% sodium dodecyl sulfate-polyacrylamide gel electrophoresis (SDS-PAGE) gels and transferred onto polyvinylidene difluoride (PVDF) membranes (Millipore, Billerica, MA, USA). For western blotting, the membranes were blocked in 5% nonfat dry milk in Tris-based saline-Tween 20 (TBS-T) at room temperature for 1 h and then incubated overnight at 4°C with various primary antibodies (Supplementary Table 3). Primary antibodies bound to the membranes were detected with horseradish peroxidase (HRP)-conjugated secondary antibodies (1:10000 dilution; Cat. #ZDR-5306, ZDR-5307, or ZSGB-BIO, Beijing, China), and the immunoblotting signal was visualized using enhanced chemiluminescence reagents (Cat. #34076; Thermo-Fisher, Waltham, MA, USA) on chemiluminescence instrument C300 (Azure, Dublin, CA, USA). The levels of individual target proteins relative to the level of the glyceraldehyde-3-phosphate dehydrogenase (GAPDH) control were determined by densitometric analysis using ImageJ software.

### Single-cell RNA sequencing (RNA-seq)

The OCT4 promoter–controlled enhanced green fluorescent protein (EGFP) expression vector was constructed by cloning the human OCT4 promoter region (1–2012 bp) using polymerase chain reaction (PCR) and inserted with a DNA fragment of EGFP cDNA into the plasmid pGL3-basic at the Nhel and BglII sites to generate pGL3-OCT4-EGFP and sequenced. Next, the CD44^−^/CD24^−^ MDA-MB-231 cells were grown and sorted by flow cytometry. The collected cells were grown and transfected with pGL3-OCT4-EGFP using Lipofectamine 3000 (Invitrogen, Carlsbad, CA, USA) and transferred into QuAscount assay plates for 1 or 3 days for capture of the single cells using a microraft (Teacon, Maennedorf, Switzerland) according to the manufacturer’s instructions at 24, 72, and 120 h and subjected to RNA-seq analysis to identify differentially expressed genes (DEGs).

The DEGs in the 24- and 72-h samples were identified by RNA-seq. In brief, total RNA was isolated from individual single-cell samples, and their mRNA was enriched using the oligo-dT microbeads and fragmented to 300–500 bp before being reversely transcribed into cDNA. The cDNA samples were amplified by PCR to generate cDNA libraries, and the latter were sequenced with an Illumina HiSeq™ (Illumina, San Diego, CA, USA). The high-quality reads were aligned to the mouse reference genome (GRCm38) using the Bowtie2 v2.4.2 (Baltimore, MD, USA), and the expression level of individual genes was normalized to the fragments per kilobase of the exon model per million mapped reads from RNA-seq by expectation maximization. DEGs were considered significant if the gene expression difference changed >2-fold compared to the control and had an adjusted p-value of <0.05.

### Bioinformatics analysis

The DEGs were further analyzed by gene ontology (GO) with the online tool AMIGO (http://www.geneontology.org) and Database for Annotation, Visualization and Integrated Discovery (DAVID) software. The enrichment degrees of differentially expressed proteins (DEPs) and DEGs were analyzed using the Kyoto Encyclopedia of Genes and Genomes (KEGG; http://www.genome.jp/kegg) annotations.

### Nude mouse tumor cell xenograft and lung metastasis assays

The experimental protocol was approved by the Animal Research and Care Committee of China Medical University (Shenyang, China) and followed the Guidelines of the Care and Use of Laboratory Animals issued by the Chinese Council on Animal Research. Female BALB/c nude mice (6 weeks old) were obtained from Human Silaikejingda Laboratory Animals (Changsha, China) and housed in a specific pathogen-free facility with free access to autoclaved food and water. CSCs from parental MDA-MB-231 and CD44^−^/CD24^−^ converted MDA-MB-231 cells were injected into individual BALB/c nude mice (2 × 10^3^ cells/mouse). Tumor growth and body weight were monitored until 21 days post-tumor cell inoculation. At the end of the experiment, subcutaneous tumors were recovered and weighed. In addition, some tumor cell xenografts were recovered from each group at 7 and 21 days after inoculation and digested to prepare single-cell suspensions for staining with FITC-anti-CD24, PE-anti-CD44, or isotype controls for flow cytometric sorting of CD44^+^/CD24^−^, CD44^−^/CD24^−^, CD44^−^/CD24^+^, and CD44^+^/CD24^+^ cells.

Moreover, an equal number (1 × 10^3^ cells/mouse) of CSCs from CD44^+^/CD24^−^ and CD44^−^/CD24^−^ converted MDA-MB-231 cells were injected into the tail vein of NOD/SCID mice and grown for 21 days to assess their lung metastasis. Afterward, the mice were sacrificed and the tumor cell lung metastasis was resected. The metastatic tissue was processed for hematoxylin and eosin (H&E) staining and photographed (n=5–8 per group). Additionally, tumor weight and sizes were measured in each group.

### Statistical analysis

The data were expressed as the mean ± the standard deviation (SD), and the differences between the groups were analyzed by chi-squared or Student’s t tests, as applicable. The statistics of all biological experiments were based on more than three independent replicated experiments. The DFS of each group of patients was estimated by the Kaplan-Meier curves and analyzed using the log-rank test. All statistical analyses were performed using SPSS 23.0 (SPSS, Inc., Chicago, IL, USA). A p-value less than or equal to 0.05 was considered statistically significant.

## Acknowledgements

This work was supported by grants from The National Natural Science Foundation of China (#81872159, #81572609, and #31601142) and the Major Project Construction Foundation of China Medical University (#2017ZDZX05).

## Author contribution

Caigang Liu conceived and designed the experiments; Xinbo Qiao, Lisha Sun and Yixiao Zhang prepared the manuscript; Qingtian Ma, Jie Yang and Liping Ai performed the experiments; Jinqi, Guanglei Chen, Hao Zhang, Ce Ji, Xi Gu, Haixin Lei,and Yongliang Yang analyzed the data.

## Supplementary information

**Figure S1.**
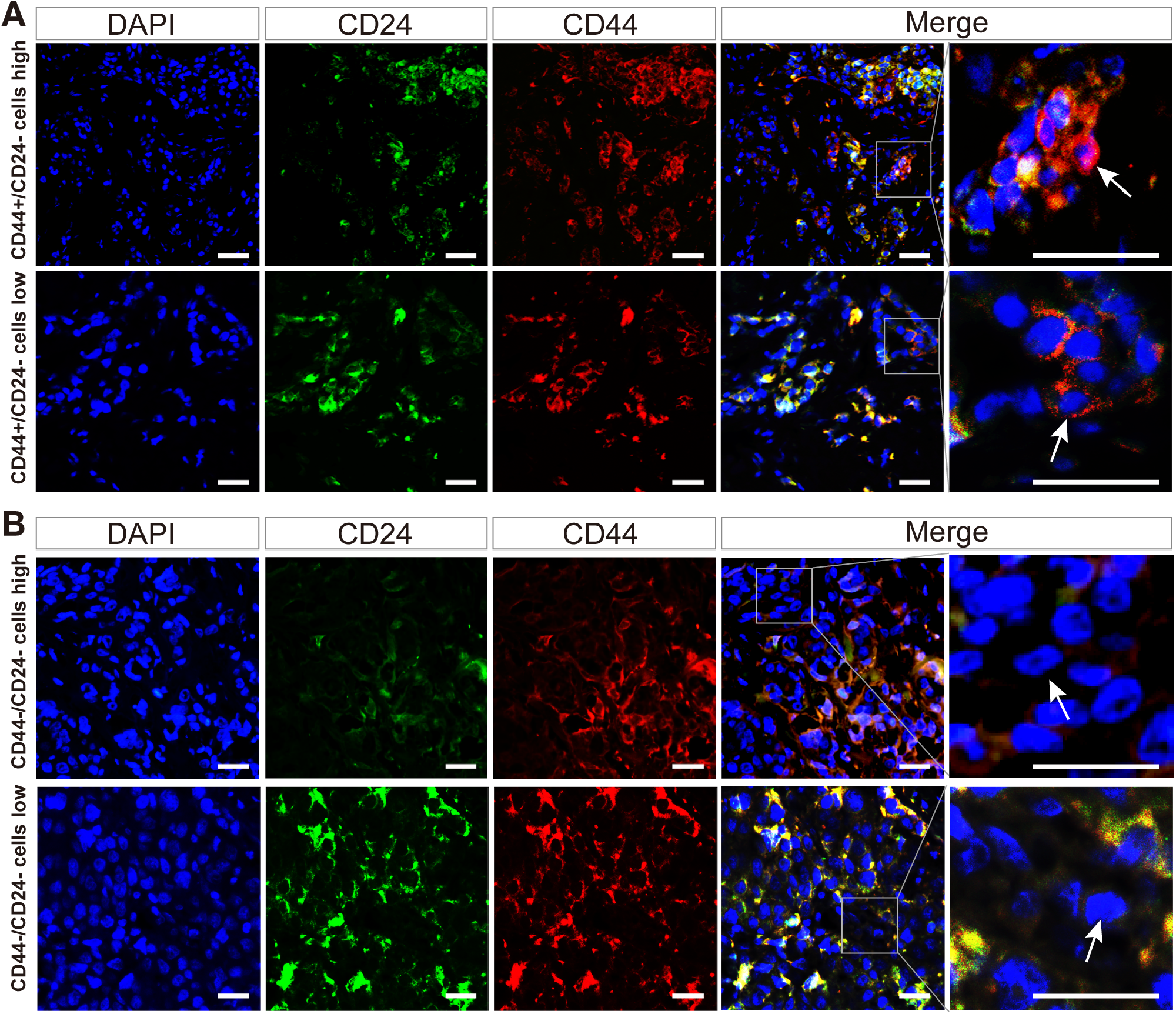
Immunofluorescent staining of CD44^+^/CD24^−^ and CD44^−^/CD24^−^ cancer cells in human breast cancer specimens. (A) Representative fluorescent images of breast cancer cells stained by anti-CD24 and anti-CD44 antibodies. Scale bar, 20 μm; high CD44^+^/CD24^−^ cells (≥2%) and low ones (<2%). (B) Representative fluorescent images of cancer cells stained by anti-CD24 and anti-CD44 antibodies. Scale bar, 20 μm; high CD44^−^/CD24^−^ cells (≥19.5%) and low ones (<19.5%).

**Figure S2.**
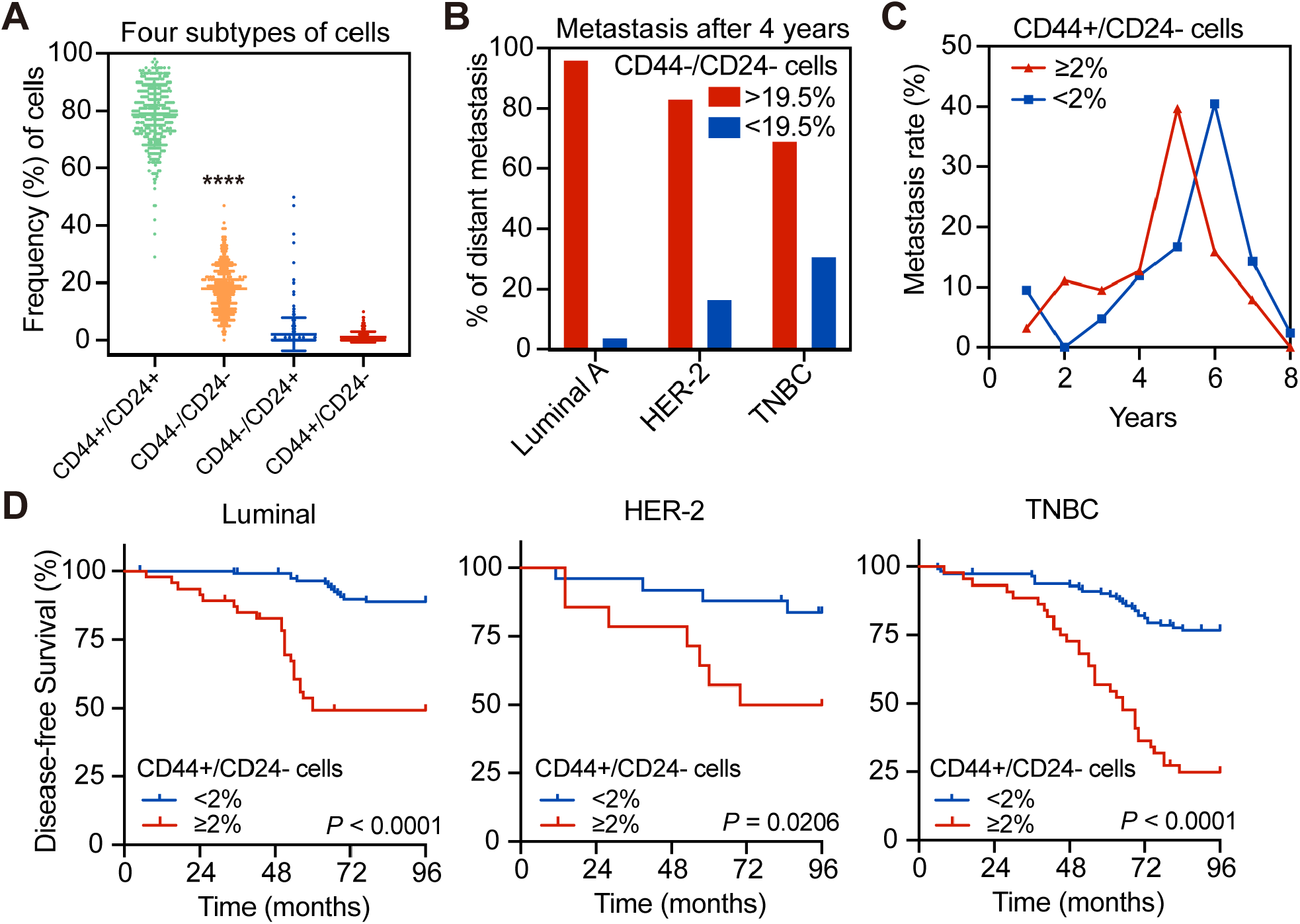
Diagnostic and prognostic value of stratification of metastatic tumors by the percentage of CD44^−^/CD24^−^ cancer cells. (A) Flow cytometric analysis of the percentages of CD44^+^/CD24^+^, CD44^−^/CD24^−^, CD44^−^/CD24^+^, and CD44^+^/CD24^−^ cells in breast cancer tissue specimens (n=355). \*\*\*\**P*<0.0001 comparing CD44^+^/CD24^−^ cells and CD44^−^/CD24^−^ cells using Student’s *t*-test. (B) Tumor metastasis in postoperative breast cancer patients after 4 years stratified by the different breast cancer molecular subtypes and the percentage of CD44^−^/CD24^−^ cells. (C) Tumor metastasis of breast cancer patients stratified by the percentage of CD44^+^/CD24^−^ cells (≥2%; n=105 vs. <2%; n=250). (D) Kaplan–Meier curves. The DFS of patients with luminal breast cancer (≥2% CD44^+^/CD24^−^ cells; n=47 vs. <2%; n = 112), HER-2-positive breast cancer (≥2% CD44^+^/CD24^−^ cells; n=14 vs. <2% one; n=25), and TNBC breast cancer (≥2% CD44^+^/CD24^−^ cells; n=44 vs. <2%; n=113).

**Figure S3.**
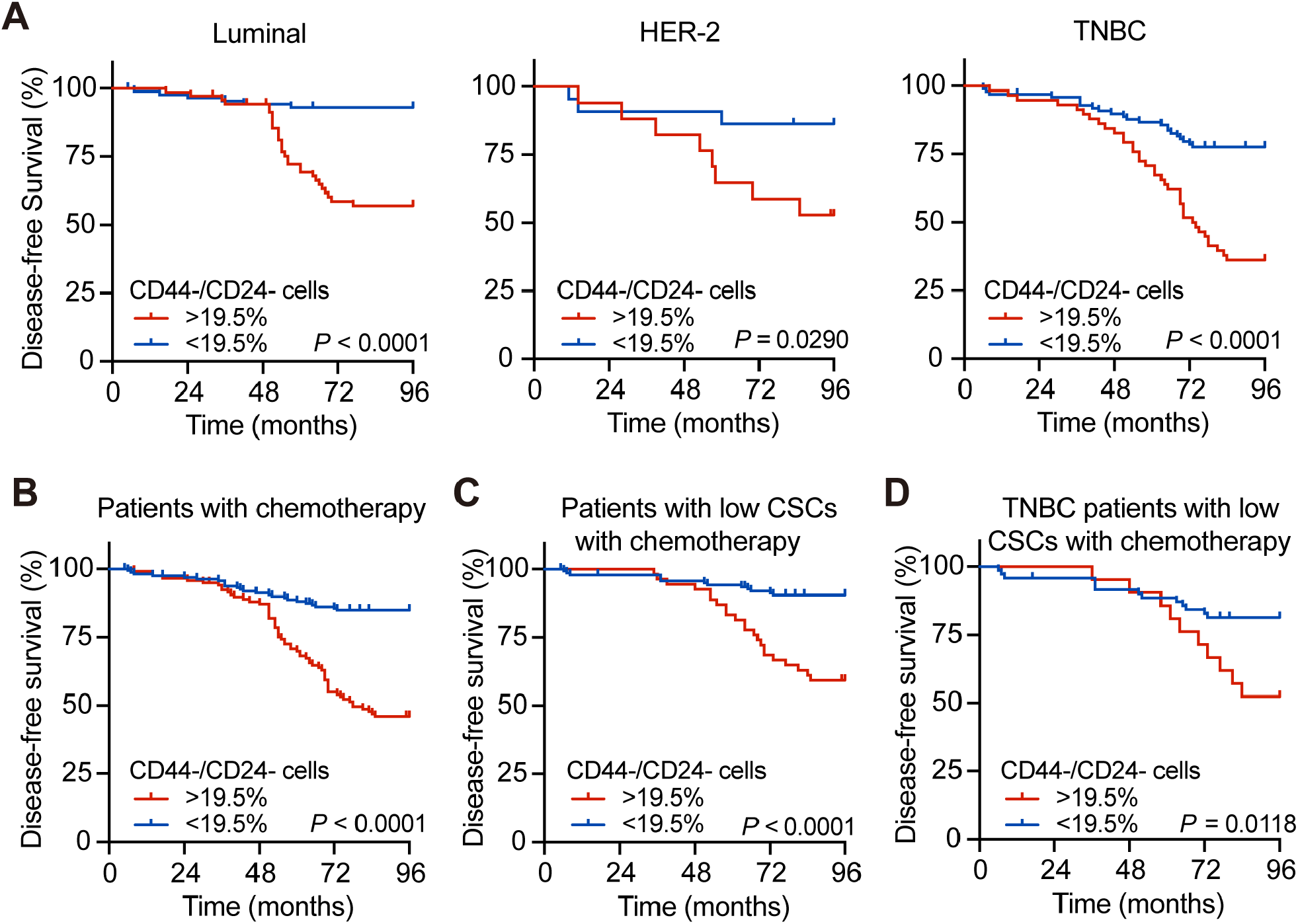
Kaplan–Meier and the log-rank test analysis of disease-free survival (DFS) in breast cancer patients stratified by percentage of CD44^−^/CD24^−^ cells. (A) Luminal (≥19.5% CD44^−^/CD24^−^ cells; n=71 vs. <19.5%; n=88), HER-2-positive (≥19.5% CD44^−^/CD24^−^ cells; n=17 vs. <19.5%; n=22), and TNBC breast cancer (≥19.5% CD44^−^/CD24^−^ cells; n=58 vs. <19.5%; n=99). (B) Stratified by chemotherapy (≥19.5% CD44^−^/CD24^−^ cells; n=118 vs. <19.5%; n=161). (C) Stratified by the low CSC level among tumor cells after chemotherapy (≥19.5% CD44^−^/CD24^−^ cells; n=54 vs. <19.5%; n=138). (D) Stratified by TNBC after chemotherapy (≥19.5% CD44^−^/CD24^−^ cells; n=21 vs. <19.5%; n=70).

**Figure S4.**
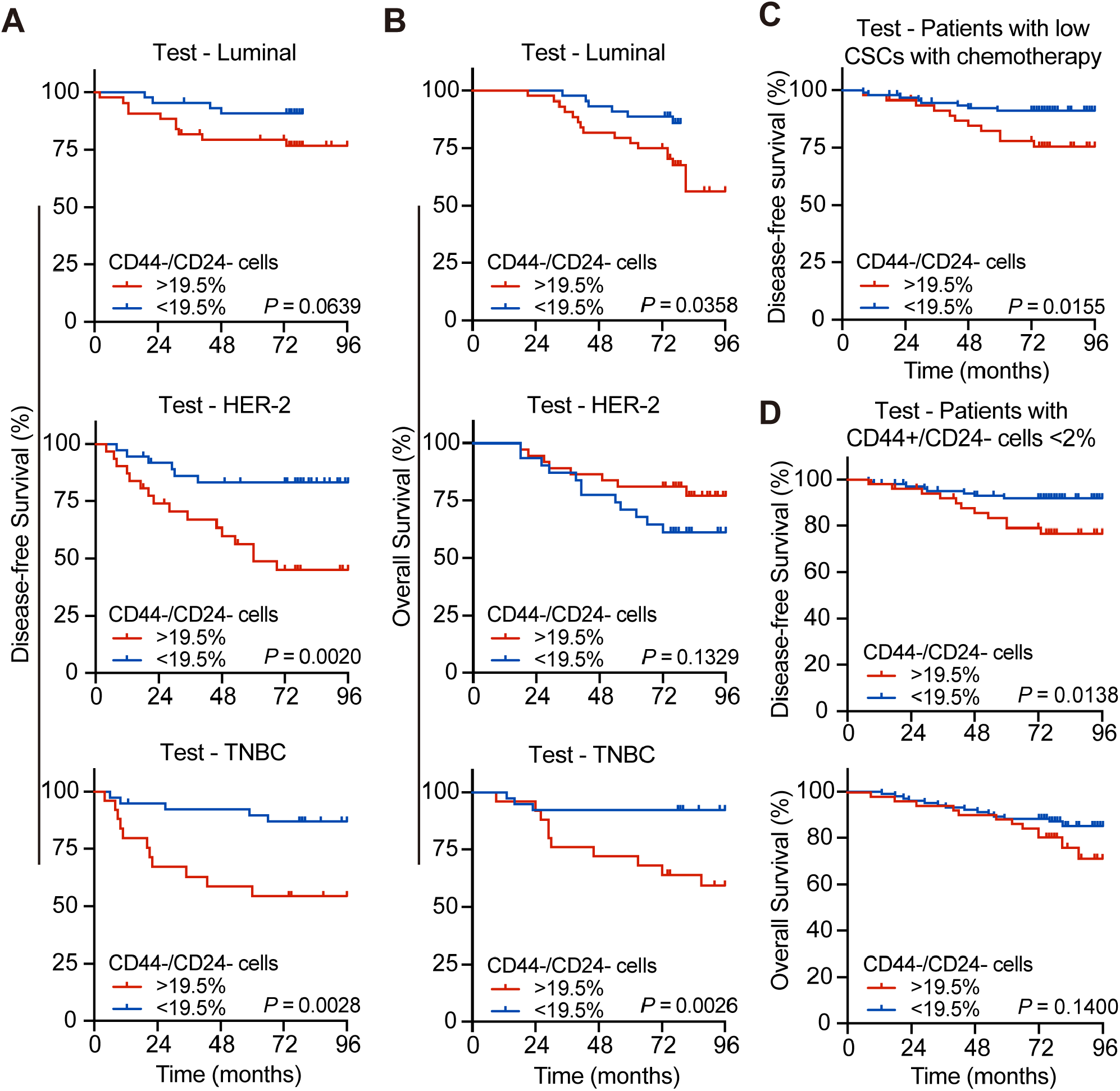
Kaplan–Meier and the log-rank test analysis of DFS and OS in the test group of patients stratified by percentage of CD44^−^/CD24^−^ cells. (A) DFS of stratified luminal cases (≥19.5% CD44^−^/CD24^−^ cells; n=44 vs. <19.5%; n=45), HER-2-positive cases (≥19.5% CD44^−^/CD24^−^ cells; n=31 vs. <19.5%; n=37), and TNBC cases (≥19.5% CD44^−^/CD24^−^ cells; n=25 vs. <19.5% ones; n=39). (B) OS of stratified luminal cases (≥19.5% CD44^−^/CD24^−^ cells; n=44 vs. <19.5%; n=45), HER-2-positive cases (≥19.5% CD44^−^/CD24^−^ cells; n=31 vs. <19.5% ones; n=37), and TNBC cases (≥19.5% CD44^−^/CD24^−^ cells; n=25 vs. <19.5%; n=39). DFS stratified by low CSC level after chemotherapy (≥19.5% CD44^−^/CD24^−^ cells; n=47 vs. <19.5%; n=94). (D) DFS and OS stratified by >19.5% CD44^−^/CD24^−^ cells (n=51) vs. < 19.5% ones (n=181) in patients with <2% CD44^+^/CD24^−^ cells.

**Figure S5.**
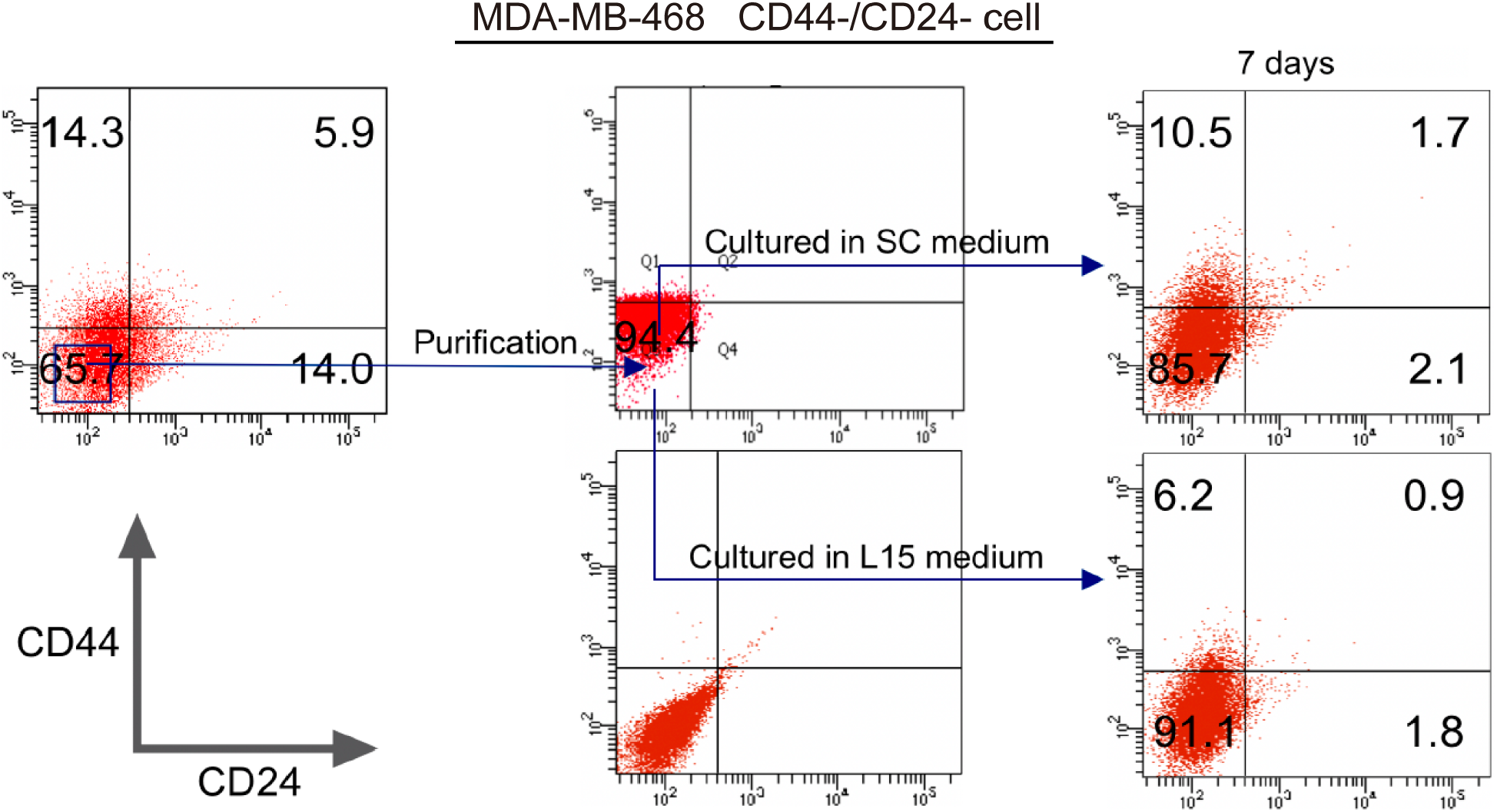
Representative images of flow cytometric data on the conversion of CD44^−^/CD24^−^ TNBC cells into CD44^+^/CD24^−^ CSCs in vitro. After sorting the CD44^−^/CD24^−^ cells by flow cytometry and culturing them in SC or L15 medium for 7 days, the percentage of CD44^+^/CD24^−^ CSCs in CD44^−^/CD24^−^ MDA-MB-468 cells was further analyzed by flow cytometry.

**Figure S6.**
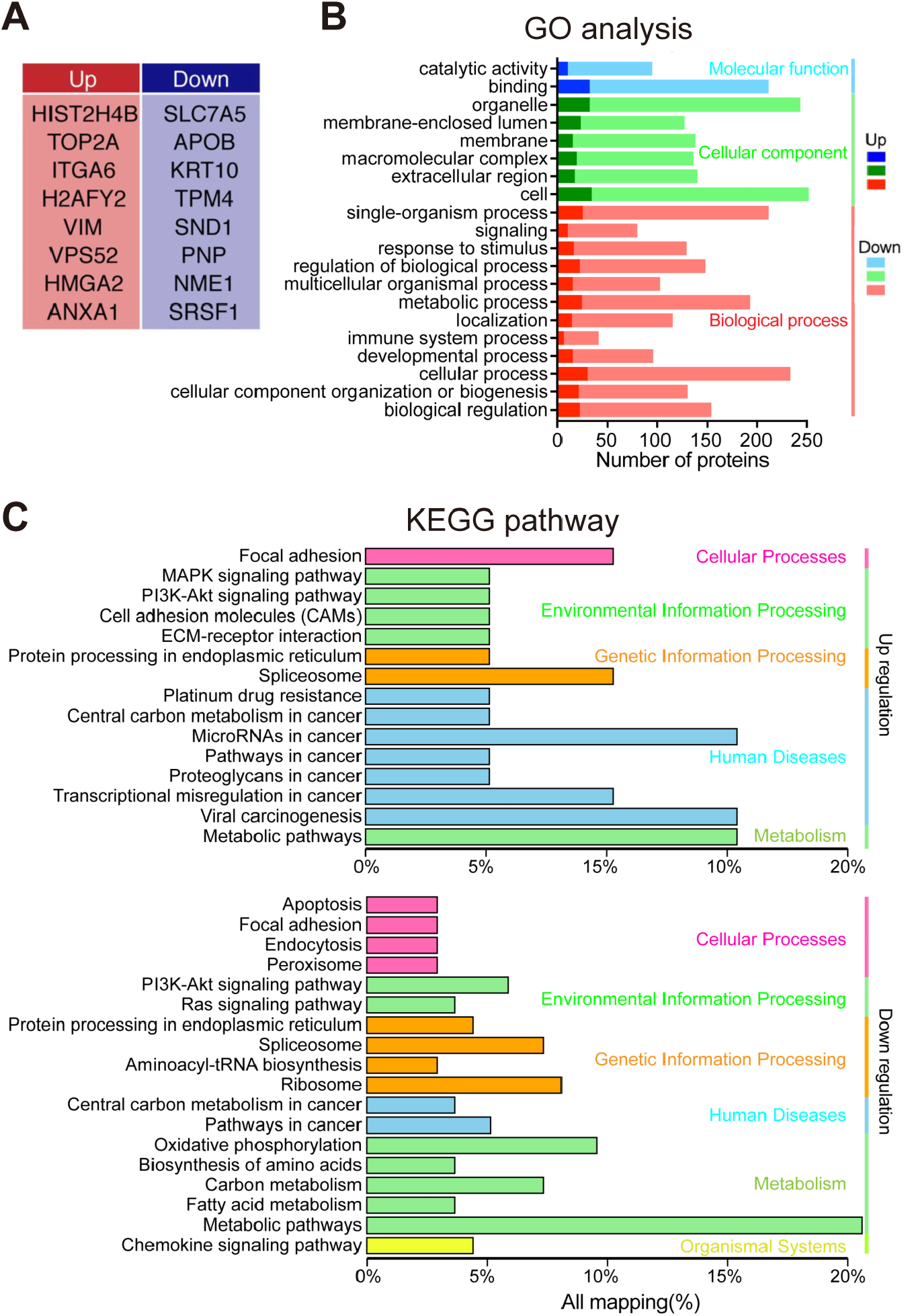
Bioinformatic analysis of DEGs between 24-h dedifferentiated single cells and CD44^−^/CD24^−^ MDA-MB-231 cells. (A) The most up- and down-regulated DEPs between 24- and 72-h samples identified by stable-isotope labeling by amino acids in cell culture assay (SILAC). (B) GO term for the DEGs. (C) KEGG analysis of the DEGs.

**Table S1.** Clinicopathological characteristics of patients

| Variables | Training set<br>n (%) | Testing set<br>n (%) |
| --- | --- | --- |
| Number of patients | 355 | 221 |
| Median age at<br>diagnosis (yrs.) | 51 | 51 |
| Molecular subtype |  |  |
| Luminal | 159 (44.79) | 89 (40.27) |
| HER-2 | 39 (10.99) | 68 (30.77) |
| TNBC | 157 (44.22) | 64 (28.96) |
| N stage |  |  |
| N0 | 240 (67.61) | 123 (55.66) |
| N1 | 69 (19.44) | 62 (28.05) |
| N2 | 25 (7.04) | 17 (7.69) |
| N3 | 21 (5.91) | 19 (8.60) |
| T stage |  |  |
| T1 | 196 (55.21) | 63 (28.51) |
| T2 | 156 (43.94) | 147 (66.51) |
| T3 | 3 (0.85) | 11 (4.98) |
| Clinical stage |  |  |
| I | 110 (30.99) | 31 (14.03) |
| II | 198 (55.77) | 156 (70.59) |
| III | 47 (13.24) | 34 (15.38) |
| Lymph node<br>involvement | 115 (32.39) | 98 (44.34) |
| Distant metastases | 105 (29.58) | 52 (23.53) |
| Bone metastases | 19 (5.35) | 22 (9.95) |
| Follow-up in months | 96 (5 - 96) | 78 (9 - 96) |
| Death | 37 (10.42) | 54 (24.43) |

**Table S2.**
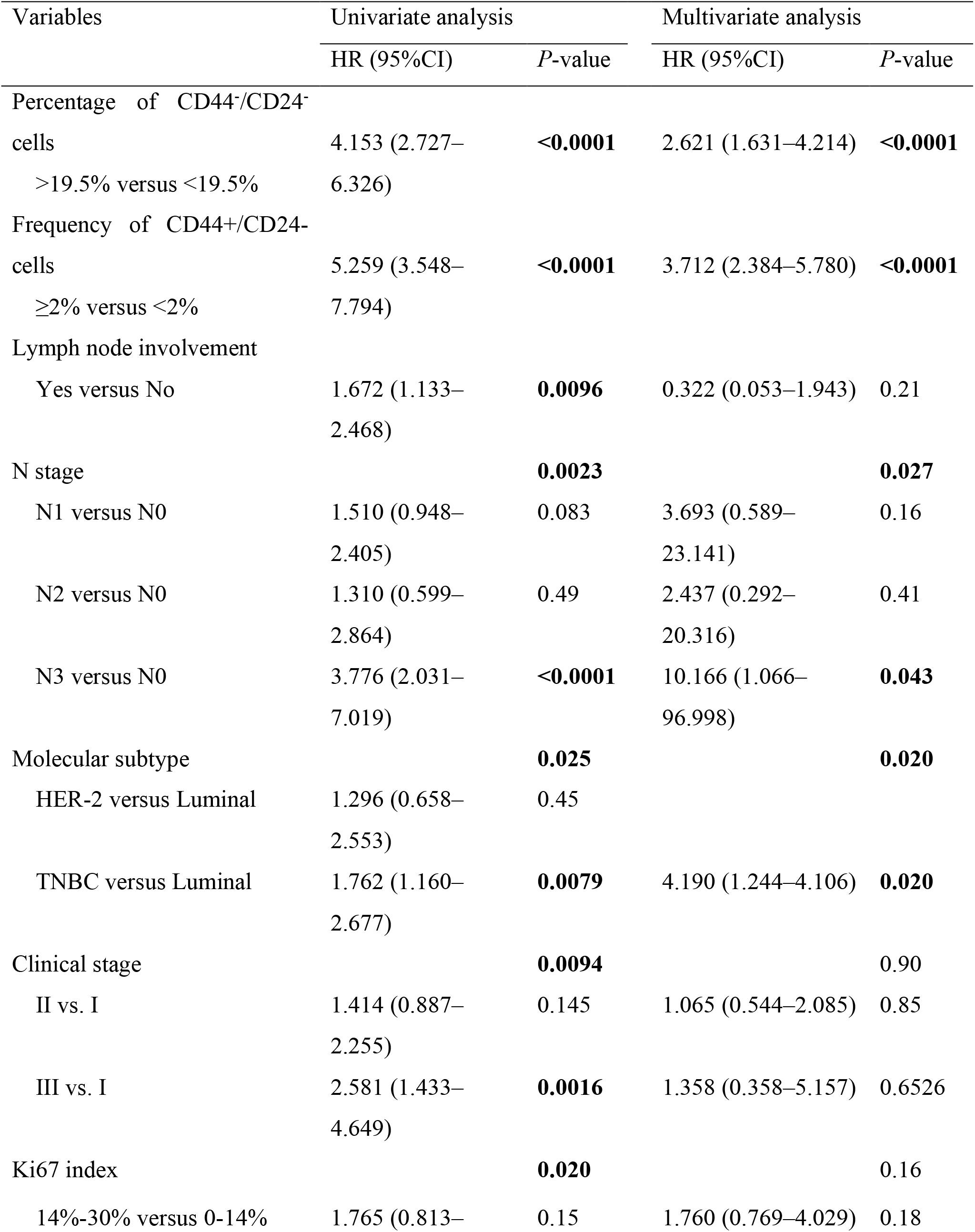
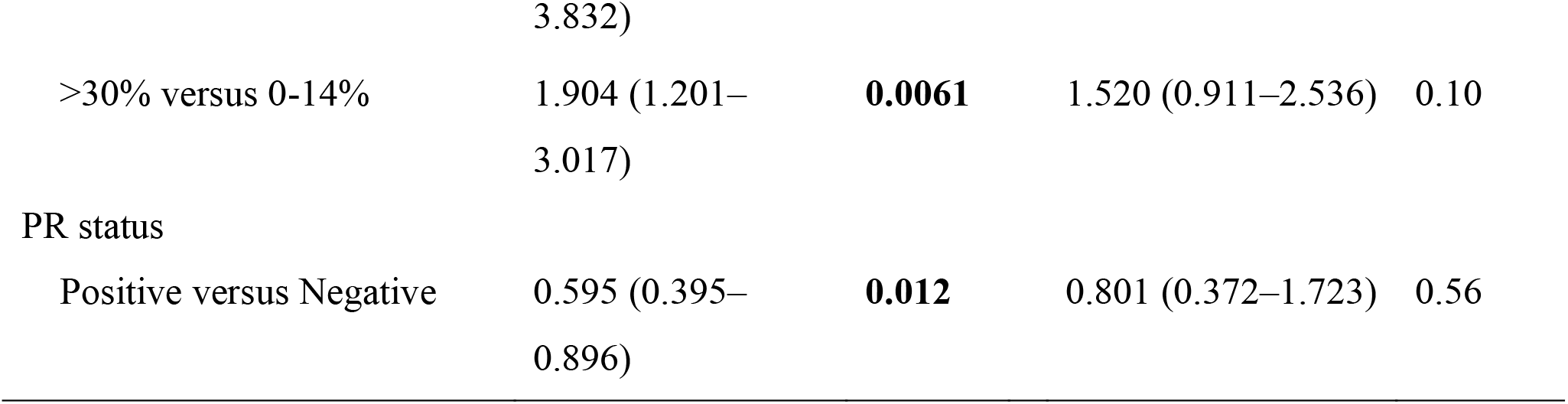
Univariate and multivariate Cox regression analyses of clinicopathological factors as predictors of DFS

**Table S3.** Primary antibodies used in western blotting

| Protein | Catalog number | Manufacturer |
| --- | --- | --- |
| GAPDH | 10494-1-AP | Proteintech (Rosemont, IL, USA) |
| OCT4 | 11263-1-AP | Proteintech |
| SOX2 | 11064-1-AP | Proteintech |
| ABCB1 | 22336-1-AP | Proteintech |
| YAP | 13584-1-AP | Proteintech |
| Tubulin-β | 10094-1-AP | Proteintech |
| Alpha-tubulin | 11224-1-AP | Proteintech |
| Phospho-YAP | KA1392C | Immunoway (Plano, TX, USA) |
| NF-κB | YM3111 | Immunoway |
| Phospho-NF-κB | YP0188 | Immunoway |
| USP31 | ab240543 | Abcam (Cambridge, MA, USA) |
| NANOG | ab80892 | Abcam |
| Nestin | ab22035 | Abcam |
| Lamin B1 | 12987-1-AP | Proteintech |

